# CRISPR/Cas9 technology abolishes the *BCR/ABL1* oncogene in chronic myeloid leukemia and restores normal hematopoiesis

**DOI:** 10.1101/2020.08.05.237610

**Authors:** Elena Vuelta, José Luis Ordoñez, Verónica Alonso-Pérez, Lucía Méndez, Patricia Hernández-Carabias, Raquel Saldaña, Julián Sevilla, Elena Sebastian, Sandra Muntión, Fermín Sánchez-Guijo, Jesús María Hernadez-Rivas, Ignacio García-Tuñón, Manuel Sánchez-Martín

## Abstract

Chronic myeloid leukemia (CML) is a hematopoietic stem cell disease produced by a unique oncogenic event involving the constitutively active tyrosine kinase (TK) *BCR/ABL1*. TK activity explains most features of CML, such as tumor development and maintenance. TK-inhibitory (TKI) drugs have changed its prognosis and natural history. Unfortunately, the *ABL1* gene persists unaffected by TKIs, leukemic stem cells (LSCs) remains, resistant mutations arise and adverse effects may occur during treatment. To address this problem, we have designed a potential therapeutic alternative with CRISPR/Cas9 genome editing nucleases that target LSCs. The strategy was successfully developed in murine and human cell lines and finally was evaluated in primary LSCs isolated from CML transgenic mice and from CML patients. Mouse CML-LSCs edited were orthotopic transplanted in immunodeficient NSG niches where restored the normal hematopoiesis. Importantly, patient-derived xenografts with CD34^+^-LSCs edited, repopulated and restored the normal hematopoiesis in immunodeficient NSG niches. We show, for the first time, how CRISPR technology efficiently interrupts the *BCR/ABL1* oncogene in murine and human LSCs to provide a significant therapeutic benefit. We propose human CML as a potential candidate for CRISPR therapy, providing proof-of-principle for genome editing in CML patients, and open new avenues for the application of this technique in other fusion genes.

**Key points:** CRISPR system destroys BCR/ABL oncogene and induces a therapeutic benefit in a CML mouse model and CML patient derived xenografts.

## INTRODUCTION

Chronic myeloid leukemia (CML) was the first neoplastic disease to be associated with a chromosomal abnormality: a reciprocal translocation between chromosomes 9 and 22 ^1,2^, fusing *BCR* and *ABL1* genes. Since then, a considerable number of fusion genes have been detected in malignant disorders, most of them closely related to the pathogenesis of specific sub-types of leukemias, lymphomas or sarcomas ^3^. In contrast to other tumors such as carcinomas, the presence of *BCR/ABL1* can explain most of the cellular features of the leukemia. Specifically, CML is a hematopoietic stem cell (HSC) disorder in which BCR-ABL1 oncoprotein has a constitutively active tyrosine kinase (TK) initiating and driving the maintenance of the disease during the chronic phase (CP) ^4–10^. The aberrant ABL1 signaling activates downstream targets that enhances cell growth, inhibits apoptosis, alters cell adhesion, produces growth-factor independence, impairs genomic surveillance and differentiation ^11–20^. This aberrant kinase signaling reprograms the cell to cause uncontrolled proliferation and results in myeloid hyperplasia and 'indolent' symptoms. Without an effective treatment, the CP evolves to an aggressive phase called the blast crisis (BC), which is characterized by the blockage of hematopoietic differentiation and accumulation of immature blast cells, resembling acute leukemia ^21,22^. In BC-CML new alterations underlay and some pathogenic mechanisms become BCR/ABL1 independent ^23^. BC triggers a bone marrow failure, and massive infiltration with immature blasts leads to patient mortality from infection, thrombosis or anemia. In CP-CML patients, the estimated 8-year survival has improved with sustained survival, low risk of progression and life expectancy is almost identical to that of the healthy population of the same age ^21,24–26^. However, despite successful outcomes in CP-CML patients, TKI resistance is observed in approximately 25% of patients, mainly due to mutations in the *BCR-ABL1* fusion gene. Other patients experience relapse after initial success ^27^. Except for a subgroup of patients who achieve a deep and sustained molecular response, TKI therapies need to be continued indefinitely because they do not completely eliminate the leukemic stem cells that remain “oncogenic-quiescent” during the treatment ^28^. Therefore, it is still necessary to seek new therapeutic alternatives, especially for TKI-resistant patients.

The recently developed clustered regularly interspaced short palindromic repeats (CRISPR)/Cas9 system, which is widely used for genome editing in all organisms ^29–31^, could be a definitive therapeutic option for these TKI-resistant patients. In this pathological cell context, the highly efficient interruption of the *BCR/ABL1* coding sequence might be an effective therapeutic option, In fact, preliminary CRISPR assays to disrupt *BCR/ABL1* have been reported, mainly in CML cell lines, with similar results on proliferation or cell survival to TKIs ^32–37^. However, unresolved critical questions about the engraft capacity, multipotency or putative therapeutic benefit of CRISPR edited LSCs are pending.

The present work demonstrates that CRISPR-Cas9 technology efficiently targets mouse and human CML cell lines and primary murine and human LSCs. We showed for the first time that CRISPR-edited LSCs colonize and repopulate the bone marrow niche of NOD-*Prkdc*^*Scid*^-*IL2rγ*^*null*^ (NSG) mice restoring the normal hematopoiesis without a myeloid bias. Patient-derived xenograft (PDX) and bone marrow transplantation assays of a CML mouse model with edited LSCs bestowed a clinical benefit. Our study provides proof-of-principle for genome editing in CML patients.

## MATERIAL AND METHODS

### *BCR/ABL1 in silico* analysis and CRISPR/Cas9 system design and reagents

In most CML patients, the point of fusion in BCR is located on the major region (M-bcr) downstream of exons 13 or 14, whereas the break-point of the ABL gene is generally located upstream of the second exon (a2) ^38,39^. This translocation produces a 210 kDa chimeric protein containing BCR as an N-terminal fusion partner joined to SH-domains, proline-rich (PxxP), nuclear localization signal (NLS), DNA-binding, nuclear export signal (NES) and actin-binding domains of ABL ^40,41^ (Supplementary figure 1).

In order to destroy the *BCR-ABL* TK domain and thereby prevent Tk activity, we designed two sgRNAs (*ABL* TK1-sgRNA and *ABL* TK2-sgRNA) (Supplementary Table 1). To easily detect and track the edited leukemic cells, both sgRNAs were designed to generate a small deletion (101 bp) targeting the *ABL* exon 6, destroying the ABL TK domain, and triggering a downstream frameshift mutation (Supplementary figure 1). TK-sgRNA sequences, were designed with the Spanish National Biotechnology Centre (CNB)-CSIC web tool BreakingCas (http://bioinfogp.cnb.csic.es/tools/breakingcas/).

#### Mouse model, cell lines, cell samples, isolation, electroporation and culture conditions

The transgenic mouse model TgP210, developed by Honda et al. ^42^, expresses the human cDNA of *BCR/ABL*p210, which is controlled by the hematopoietic stem cell *Tec* promoter, mimicking human CML disease.

Boff-p210 is a murine interleukin 3 (IL3)-independent cell line derived from the hematopoietic cell line Baf/3^43^ that expresses the *BCR/ABL* cDNA transgene ^44–46^. Boff-p210 was maintained in Dulbecco’s Modified Eagle’s Medium (DMEM) (Gibco, Thermo Fisher Scientific, CA, USA) supplemented with 10% FBS and 1% penicillin/streptomycin (Thermo Fisher Scientific, CA, USA). IL-3-dependent Baf/3 cells, used as a parental cell line, were grown in the same medium supplemented with 20% WEHI-3-conditioned medium as a source of IL-3.

The human CML-derived cell line K562 was purchased from the DMSZ collection (Leibniz-Institut DSMZ-Deutsche Sammlung von Mikroorganismen und Zellkulturen GmbH, Germany). K562 cells were cultured in RPMI 1640 medium (Gibco, Thermo Fisher Scientific, CA, USA) supplemented with 10% FBS, and 1% penicillin/streptomycin (Thermo Fisher Scientific, CA, USA). All cell lines were incubated at 37°C in a 5% CO_2_ atmosphere. The presence of mycoplasma was tested for frequently in all cell lines with a MycoAlert kit (Lonza, Switzerland), exclusively using mycoplasma-free cells in all the experiments carried out.

Mouse hematopoietic cells were isolated from mouse bone marrow samples by flushing their femurs with a syringe containing 2 mL PBS–10% FBS. After red blood lysis treatment, AutoMACs sorter with a Direct Lineage Cell Depletion Kit mouse (MiltenyiBiotech, Germany) was used to separate Lin^−^ stem cells, which were then maintained in culture for 24 h in IMDM (Gibco, Thermo Fisher Scientific, CA, USA) supplemented with 2% fetal bovine serum, interleukin-3 (mIL-3; 10 μg/mL), interleukin-6 (mIL-6; 10 μg/mL), and thrombopoietin (mTPO; 10 μg/mL) and stem cell growth factor (mSCF; 10 μg/mL) (Preprotech EC Ltd., UK). Mice were housed in a temperature-controlled specific pathogen-free (spf) facility using individually ventilated cages, standard diet and a 12 h light/dark cycle, according to EU laws, at the Servicio de Experimentación Animal at the University of Salamanca.

Human CD34^+^ cells were isolated from patient bone marrow samples by erythrocyte lysis buffer treatment, followed by separation with AutoMACs sorter and CD34 Microbead Kit human (MiltenyiBiotech, Germany) and grown in IMDM (Gibco, Thermo Fisher Scientific, CA, USA) supplemented with 2% FBS (Gibco, Thermo Fisher Scientific, CA, USA), interleukin-3 (hIL-3; 10 μg/mL), interleukin-6 (hIL-6; 10 μg/mL), and thrombopoietin (hTPO) (10 μg/mL), stem cell growth factor (hSCF; 10 μg/mL) and Fms-related tyrosine kinase 3 ligand (hFlt3-L; 10 μg/mL) (Preprotech EC Ltd, UK). CD34^+^ cells were maintained in culture for 24 h before electroporation.

#### CRISPR/Cas9 ribonucleocomplex assembly and electroporation

The guide RNAs against *ABL* TK domains were composed by annealing equimolar concentrations of crRNA (specific target sequence; Integrated DNA Technology, Belgium) and of tracrRNA to a final duplex concentration of 44 μM by heating at 95°C for 5 min and ramp down temperature to 25°C. 22 pmol of duplex was incubated with 18 pmol of Cas9 enzyme (Integrated DNA Technology, Belgium) to final volume of 1 μL for electroporation. We added 2 μL of 10.8 μM of Electroporation Enhancer (Integrated DNA Technology, Belgium), and 9 μL of cell suspension of 3 × 10^7^ cells/mL.

Cells were electroporated using Neon Transfection System 10 μL Kit (Invitrogen, Thermo Fisher Scientific, CA, USA) following the manufacturer’s instructions. The electroporation parameters for each cell type are described in the following table:

| Cells | Pulse Voltage (v) | Pulse Width (ms) | Pulse Number |
| --- | --- | --- | --- |
| Boff-p210 | 1700 | 2 | 1 |
| K562 | 1450 | 10 | 3 |
| mLin<sup>-</sup> | 1700 | 2 | 1 |
| hCD34<sup>+</sup> | 1600 | 10 | 3 |

#### DNA and RNA insolation, amplification and genotyping

Genomic DNA from cells was extracted using the QIAamp DNA Micro Kit (Qiagen) following the manufacturer’s protocol. To amplify the different target regions of human ABL-1, PCR was performed with the oligos described in Supplementary table 1.

PCR products were purified using a High Pure PCR Product Purification Kit (Roche) and sequenced by the Sanger method using forward and reverse PCR primers.

#### RT-PCR and quantitative PCR

To test BCR/ABL mRNA expression, 150 ng of total mRNA from cells were extracted by RNeasy mini kit (Quiagen, Germany) and *in vitro*-retrotranscribed using a SuperScript III First-Strand Synthesis Super Mix kit (Thermo Fisher Scientific, CA, USA). cDNA was used as a template to PCR-amplify the specific edited sequence of *BCR/ABL* exon 6 (Supplementary table 1). PCR product was purified and Sanger-sequenced.

BCR/ABL mRNA expression was quantified by qPCR (Applied Biosystems, CA, USA) using oligos targeting *BCR/ABL* Phusion point BCR/ABL qPCR F and BCR/ABL qPCR R. GAPDH mRNA expression was used as a control (oligos Gapdh qPCR F and R, Supplementary table 1).

#### Western blot and immunofluorescence

BCR-ABL protein expression was assessed by SDS-PAGE and western blotting using a mouse anti-ABL antibody (1:1,000, Santa Cruz Biotechnology, CA, USA). Horseradish peroxidase-conjugated α-mouse antibody (1:10,000; NA931V, GE Healthcare, IL, USA) was used as a secondary antibody. Antibodies were detected using ECL™ Western Blotting Detection Reagents (RPN2209, GE Healthcare, IL, USA). As a control, β-actin expression was measured using mouse anti-β-actin (1:20,000; Sigma Aldrich, USA).

For immunofluorescence analysis, cells were fixed and permeabilized onto poly-L lysine-coated slides as previously described ^47^. After blockade, slides were incubated with anti-ABL (Santa Cruz Biotechnology, CA, USA) antibodies at 1:1,000 dilution for 2 h. Cy™5 Goat Anti-Mouse (Jackson Immunoresearch, PA, USA) were used as secondary antibodies (1:1,000; 1 h). Nuclei were stained with DAPI (4′,6′-diamidine-2-fenilindol), diluted in PBS and incubated for 3 min with agitation at room temperature. Cells were washed twice in PBS and slides were mounted with Vectashield reagent (Vector Laboratories, CA, USA). Images were acquired using a Leica TCS SP5DMI-6000B confocal microscope (Leica Biosystems, Germany).

#### Apoptosis and cell-cycle analysis

Apoptosis was measured by flow cytometry with an annexin V-Dy634 apoptosis detection kit (ANXVVKDY, Immunostep, Spain) following the manufacturer’s instructions. Briefly, 5 × 10^5^ cells were collected and washed twice in PBS, and labeled with annexin V-DY-634 and non-vital dye propidium iodide (PI), to enable the discrimination of living cells (annexin-negative, PI-negative), early apoptotic cells (annexin-positive, PI-negative) and late apoptotic or necrotic cells (annexin-positive, PI-positive). In parallel, cell distribution in the cell cycle phase was also analyzed by measuring DNA content (PI labeling after cell permeabilization). Data were analyzed using FlowJo software (vX.0.7. TreeStar, OR, USA).

#### Flow cytometry analysis and cell sorting for Cas9-mediated editing and for isolating single edited cell-derived clones

Boff-p210 cells were selected by fluorescence-activated cell sorting (FACS) using FACS Aria (BD Biosciences, CA, USA) 48 h after electroporation with a CRISPR/Cas9 system composed of two *ABL*-TKs crRNA and fluorescent ATTO-labeled tracRNA (Integrated DNA Technology, Belgium). Single cells were seeded in a 96-well plate establishing the SC-*ABL*-TK and SC-Cas9 clones, the latter being used as a control.

### Hematopoietic stem cell transplantation

After 24 h in cytokine enriched culture, 1.8 × 10^6^ Lin^−^ cells from Tgp210^+^ C57BL/6 mice, or human CD34^+^ cells from the CML bone marrow sample, were divided to undergo only Cas9 (used as control) or full CRISPR system (Cas9 and *ABL*-TK sgRNAs) electroporation.

As a control of CML disease and normal hematopoiesis, Tgp210^+^ and C57BL/6J wild type mouse were used. In human-mouse bone marrow transplantation, we used 9 × 10^5^ hCD34^+^ cells without electroporation from CML patients and healthy donors as controls.

Electroporated and non-electroporated hematopoietic stem cells (HSC) were injected intravenously through the lateral tail vein in sublethally irradiated NSG (NOD scid gamma, Charles River) 4-5-week-old mice (2 Gy, 4 h before injection). Mice were irradiated at the Radioactive Isotopes and Radioprotection service of NUCLEUS, University of Salamanca. Mice were sacrificed by anesthetic overdose 4 or 6 months after cell injection.

### Flow cytometry analysis

Four months after mouse-mouse bone marrow transplantation, recipient mice were euthanized and flow cytometric analysis of peripheral blood was carried out. Red blood cells were lysed with erythrocyte lysis buffer, and the remaining cells were washed twice in PBS. Samples were stained with fluorophore-conjugated antibodies against anti-mCD45 (PerCP-Cy5), anti-mB220 (PE), anti-mIgM (APC), anti-mCD4 (FITC), anti-mCD8 (APC) (all from Biolegend, CA, USA), and against anti-Gr1 (APC-Cy7), anti-Mac (FITC), anti-sca1 (PE) and anti-ckit (PE-Cy7 (all from BD Biosciences, CA, USA).

In human-mouse bone marrow transplantation, recipient mice were euthanized 24 weeks after cell injection. Bone marrow was extracted by flushing tibia and femurs, and red cells were lysed. The remaining cells were stained with fluorophore-conjugated antibodies against anti-hCD45 (FITC), anti-hCD34 (APC), anti-hCD14 (APC-H7), anti-hCD117, anti-hCD15 (FITC) (all from BD Biosciences, CA, USA), anti-mCD45 (PerCP-Cy5, Biolegend CA, USA), and anti-hCD19 (PE-Cy7, Immunostep, Spain). Samples were obtained on a FACSAria flow cytometer (BD Biosciences CA, USA) and data were analyzed using FlowJo software (TreeStar, OR, USA).

#### Statistical analysis

Statistical analyses were performed using GraphPad Prism 6 Software (GraphPad Software). Group differences between levels of annexin V labeling and BCR/ABL expression were tested with one- and two-way ANOVA, respectively, and Tukey’s multiple comparisons test. Differences in percentages of hematological populations were estimated by two-way ANOVA. Statistical significance was concluded for values of p < 0.05 (**) and p < 0.001 (***).

#### Ethics statement

The study with human samples and human data followed the Spanish Biomedical Research Law 14/2007, RD 1716/2011, RD 1720/2007 and European Regulation 2016/679 (General Data Protection Regulation). The study was approved by the Ethics Committee for research in human drugs (CEIC) of IBSAL, Salamanca, Spain (reference PI5505/2017). The study with animals followed Spanish and European Union guidelines for animal experimentation (RD 1201/05, RD 53/2013 and 86/609/CEE) and was approved by the Bioethics Committee of the University of Salamanca and Conserjería de Agricultura y Ganadería de la Junta de Castilla y León (registration number 359).

## RESULTS

### 1. The CRISPR/Cas9 deletion system efficiently disrupts the BCR/ABL oncogene, preventing its expression

The murine CML cell line Boff-p210 was electroporated with Cas9 RNP joined to two specific sgRNAs targeting the *ABL* TK domain at exon 6 (here named Boff-*ABL*-TK) or without sgRNAs as a control (Boff-Cas9). A 473-bp band corresponding to BCR/ABL genomic exon 6 was amplified by PCR in Boff-*ABL*-TK and Boff-Cas9 cells. As expected, an extra band of 372 bp was detected only in Boff-*ABL*-TK cells, suggesting the presence of a deletion generated by the action of both RNA guides (Figure 1A). Sanger sequencing revealed a mixture of sequences between the expected cleavage point of both guides (Figure 1B) and confirmed the specific deletion at expected Cas9 cut sites. To analyze the effect of this deletion, we made a single-cell seeding (SC) and isolated a single-cell-derived clone carrying the specific CRISPR deletion affecting the ABL1 TK domain (named SC-*ABL*-TK). PCR amplification of *ABL* exon 6 showed a single band of 368 bp in SC-*ABL*-TK cells, in contrast with a 469-bp unedited band observed in SC-Cas9 cells, suggesting the presence of a specific CRISPR deletion in *ABL* loci (Figure 1A). The specific 101-bp deletion was confirmed by Sanger sequencing in Boff-*ABL*-TK and SC-*ABL*-TK (Figure 1B). To determine the relevance of the 101-bp DNA deletion at the mRNA and protein levels, ABL expression was quantified by RT-PCR. As expected, SC-*ABL*-TK cells showed a shorter human ABL mRNA than full-length sequences (Figure 1C). Sanger sequencing of the RT-PCR products showed a mRNA carrying a 101-bp deletion in the edited SC-*ABL*-TK cells. *In silico* analysis of the truncated ABL mRNA showed a 101-bp deletion that gives rise to a frameshift mutation due to a premature STOP codon at the contiguous *ABL* exon 7 (Figure 1C). To determine whether the frameshift mutation triggers mRNA decay, a qPCR was performed using oligos located downstream from the deletion region (the exon 7-8 junction; Supplementary table 1). SC-*ABL*-TK cells showed a statistically significant lower level of expression (2.6%, p<0.001) compared with SC-Cas9 control cells (Figure 1D). These data were confirmed by western blot using an ABL antibody to detect human ABL protein expression in cells. While Boff-Cas9 and SC-Cas9 cells normally expressed a 250-kDa BCR/ABL protein, SC-*ABL*-TK edited cells showed no expression of BCR/ABL protein similar to *BCR/ABL*-negative BaF/3 cells (Figure 1E).

**Figure 1.**
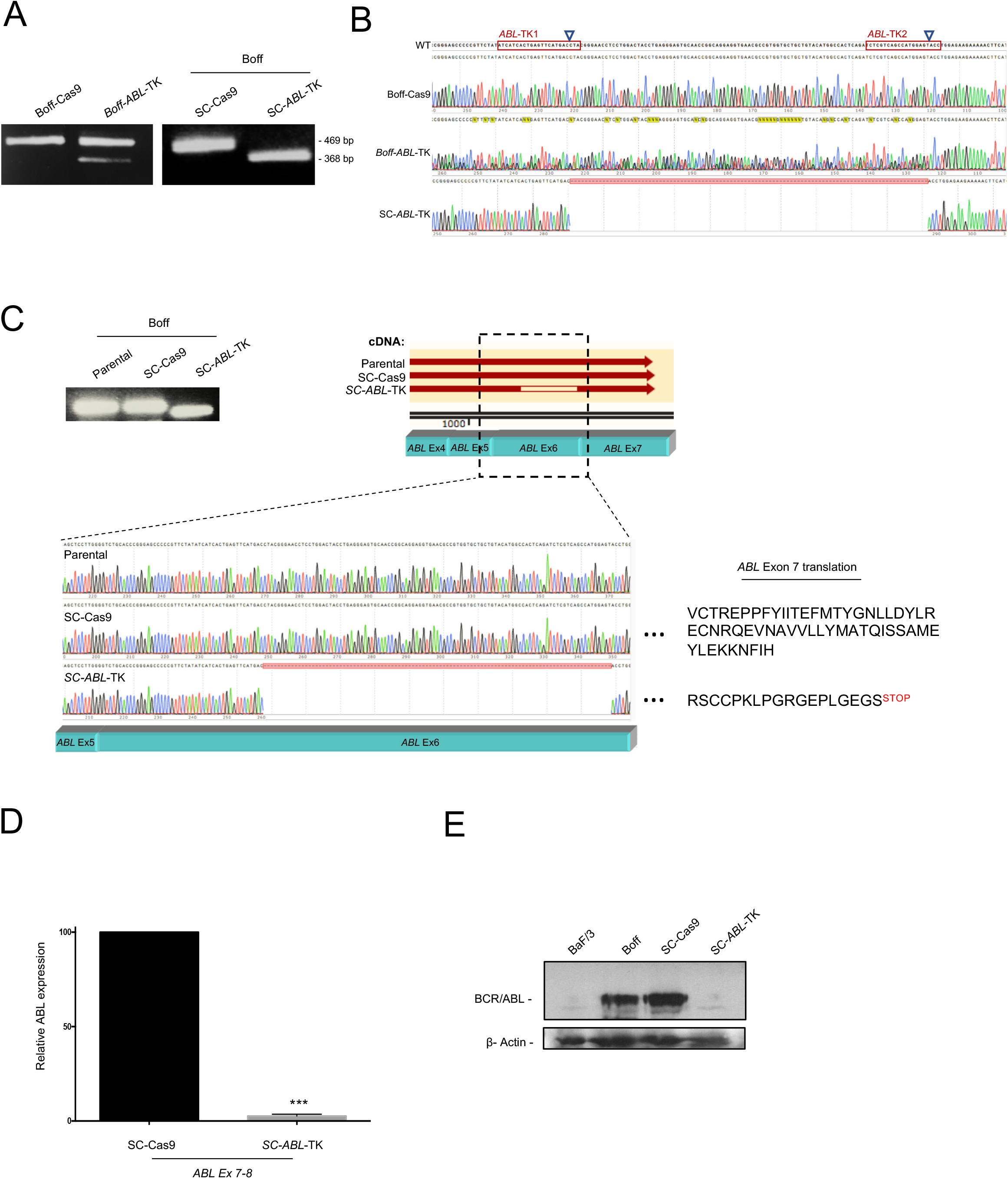
Functional analysis of the CRISPR/Cas9 deletion system in the Boff-p210 cell line. (A) PCR amplification of ABL exon 6 in Boff-p210 cells electroporated with solely Cas9 (Boff-Cas9), with Cas9 and both sgRNAs (Boff-*ABL*-TK), and in single-cell clones derived (SC-Cas9 and SC-*ABL*-TK). (B) Sanger sequencing of the amplified region of *ABL* exon 6. (C) RT-PCR of *ABL* exon 6 showing a smaller mRNA transcript in SC-*ABL*-TK than in control cells (parental and SC-Cas9). The 101-bp deletion was confirmed by Sanger sequencing in SC-*ABL*-TKs. *In silico* analysis shows this deletion generates a premature stop codon in *ABL* exon 7. (D) Quantitative PCR of ABL expression in Boff-p210 single-edited cell-derived clones (SC) at the exon 7-8 level. (mean ± SEM; *** p<0.001). (E) Western blot of BCR-ABL in Baf/3, Boff, and SC cells.

To establish whether these results are also reproducible in human cells, the human CML-derived cell line K562 was electroporated with Cas9 RNP joined to two TK sgRNAs targeting *BCR/ABL* exon 6. A 473-bp band corresponding to *BCR/ABL* genomic exon 6 was amplified by PCR in K562-Cas9 cells (control) and K562-*ABL*-TK cells. As expected, an extra band of 372 bp was amplified in the edited cells, which was consistent with CRISPR-induced deletion (Figure 2A). Sanger sequencing showed a mixture of sequences between the expected cleavage points of both guides (Figure 2A) and confirmed the specific deletion at the expected Cas9 cut sites. *BCR/ABL* oncogene expression was determined by qPCR (Figure 2B), which revealed significantly (p<0.001) lower BCR/ABL mRNA levels in K562 *ABL*-TK cells than in the two control cell lines (parental and Cas9 cells). Western blot analysis of ABL and BCR-ABL proteins in K562 *ABL*-TK cells confirmed the lower level of expression of both proteins compared with the K562-Cas9 control cells (Figure 2C). The ABL immunofluorescence assay detected a huge decrease in BCR/ABL-positive cells in the K562 *ABL*-TK cell pool (Figure 2C).

**Figure 2.**
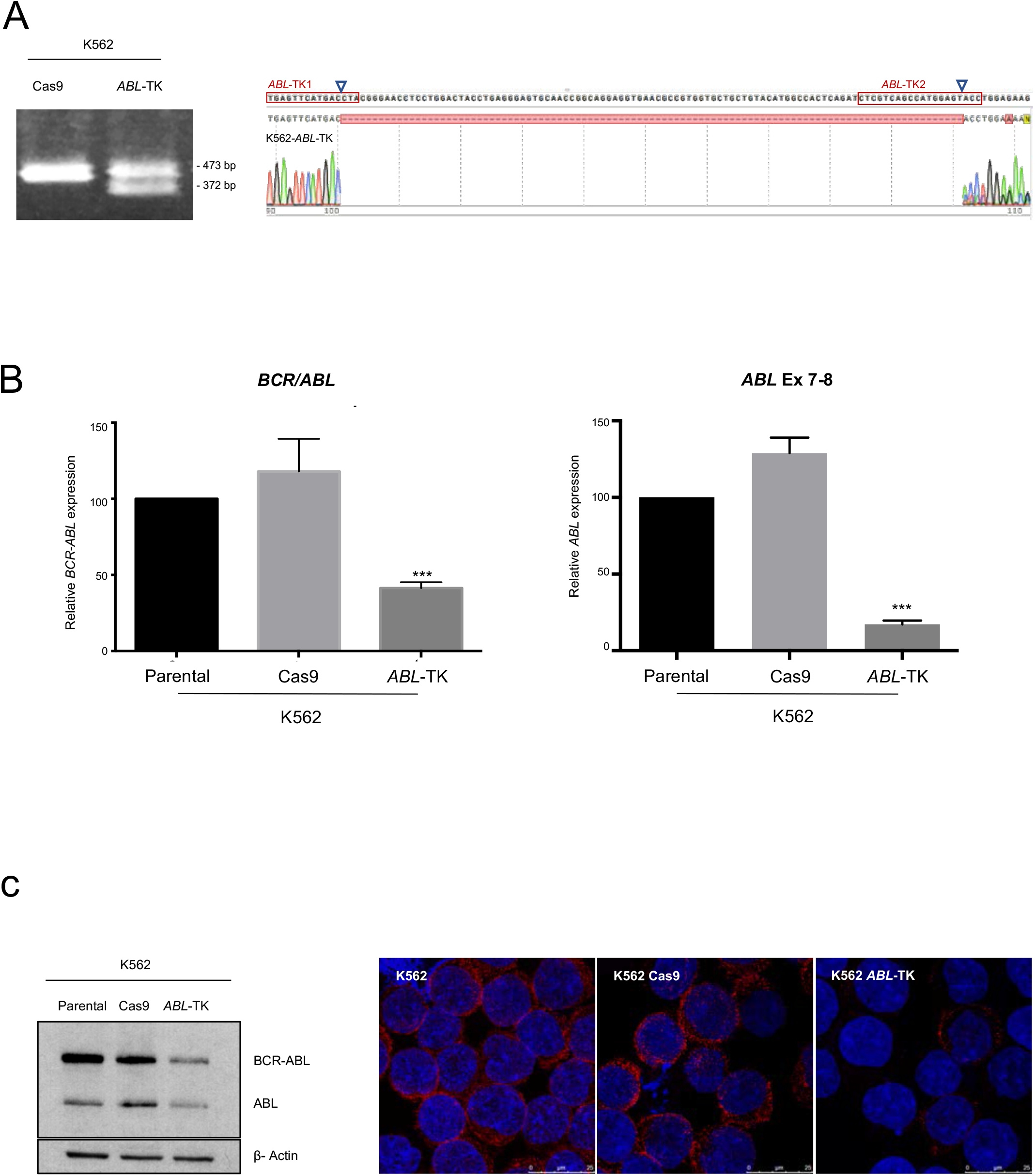
Functional analysis of the CRISPR/Cas9 deletion system in the K562 human cell line. (A) PCR amplification of ABL exon 6 in K562 cells electroporated with Cas9 nuclease (K562-Cas9) and with Cas9 joined to both sgRNAs (K562-*ABL*-TKs). Sanger sequencing of the amplified region revealed a 101-bp deletion between the expected cleavage points. (B) Quantitative PCR of *BCR/ABL* (left) and *ABL* (right) mRNA in K562 (parental), K562-Cas9 and K562-*ABL*-TK (mean ± SEM; *** p<0.001). (C) Western blot of ABL protein in parental, K562-Cas9 and K562-*ABL*-TK (left). These results were corroborated by immunocytochemistry (right).

### 2. The CRISPR/Cas9 deletion system abolishes the BCR/ABL oncogenic effect in murine and human CML cell lines

To determine the biological significance of disrupting the coding sequence of *BCR/ABL* with the CRISPR system at ABL1 TK domain, we analyzed the apoptosis levels and DNA content of edited and control cells. Unlike BaF3 (Boff parental cells), Boff-p210 cells are IL-3-independent of growth and survival due to the oncogene signaling. Thus, we studied the effect of abrogating *BCR/ABL* by CRISPR in Boff-p210 cells in the absence of IL-3 (Supplementary figure 2). 48 hours after electroporation and IL-3 withdrawal, we observed a basal level of mortality of 15.1% (annexin V-positive cells) similar to the observed in IL3 presence (13.3%) in SC-Cas9, while this level reached 99.1% on average in the SC-ABL-TK cells (Supplementary figure 2A). Accordingly, we also observed a 94.5% of SubG0 DNA content cells in SC-*ABL*-TK with respect to unedited SC-Cas9 cells (2.1%) (Supplementary figure 2B).

For the human cell line, we analyzed the level of apoptosis and the DNA content in human K562-Cas9-TK cells (Supplementary Figure 3). We also detected a significant (p<0.001) increase in the percentage of annexin V-positive cells (63.2%) in K562 *ABL*-TK-edited cells, while K562 (parental) and K562-Cas9 cells showed lower levels (8.4% and 14.5%, respectively) (Supplementary figure 3A, 3B). DNA content analysis (propidium iodide) of K562-Cas9-TKs cells showed 24.6% of SubG0 cells, while K562 (parental) and K562-Cas9 control cells showed 1.76% and 4.87% of SubG0 cells, respectively (Supplementary figure 3C).

### 3. CRISPR/Cas9 deletion system abolishes BCR/ABL oncogene expression in primary murine CML leukemic stem cells, which restore their own multipotent capacity and impair the myeloid bias

To address whether a primary murine CML stem cell, edited with the CRISPR/Cas9 system to abolish ABL1 expression, re-acquires its physiological multipotency, we used a CML transgenic mouse expressing *BCR/ABL1* human oncogene ^42^. Lin^−^ primary leukemic stem cells (mLSCs) were isolated from CML transgenic mice with symptomatic chronic disease (>60% Gr1^+^ cells in peripheral blood). *BCR/ABL* transgene sequence was disrupted by electroporation of Cas9 RNP joined to *ABL-*TK sgRNAs targeting the TK domain at exon 6. PCR amplification of *ABL* exon 6 at mLSCs-*ABL*-TK cells showed specific 469-bp and 368-bp bands, while a single band (469 bp) was observed in mLSCs control cells (Figure 3A). Sanger sequencing displayed a mixture of sequences at the CRISPR-Cas9 target region in mLSCs-*ABL*-TK cells (Figure 3A). PCR-cloning and sequencing demonstrated the specific 101-bp deletion between Cas9 cleavage points.

**Figure 3.**
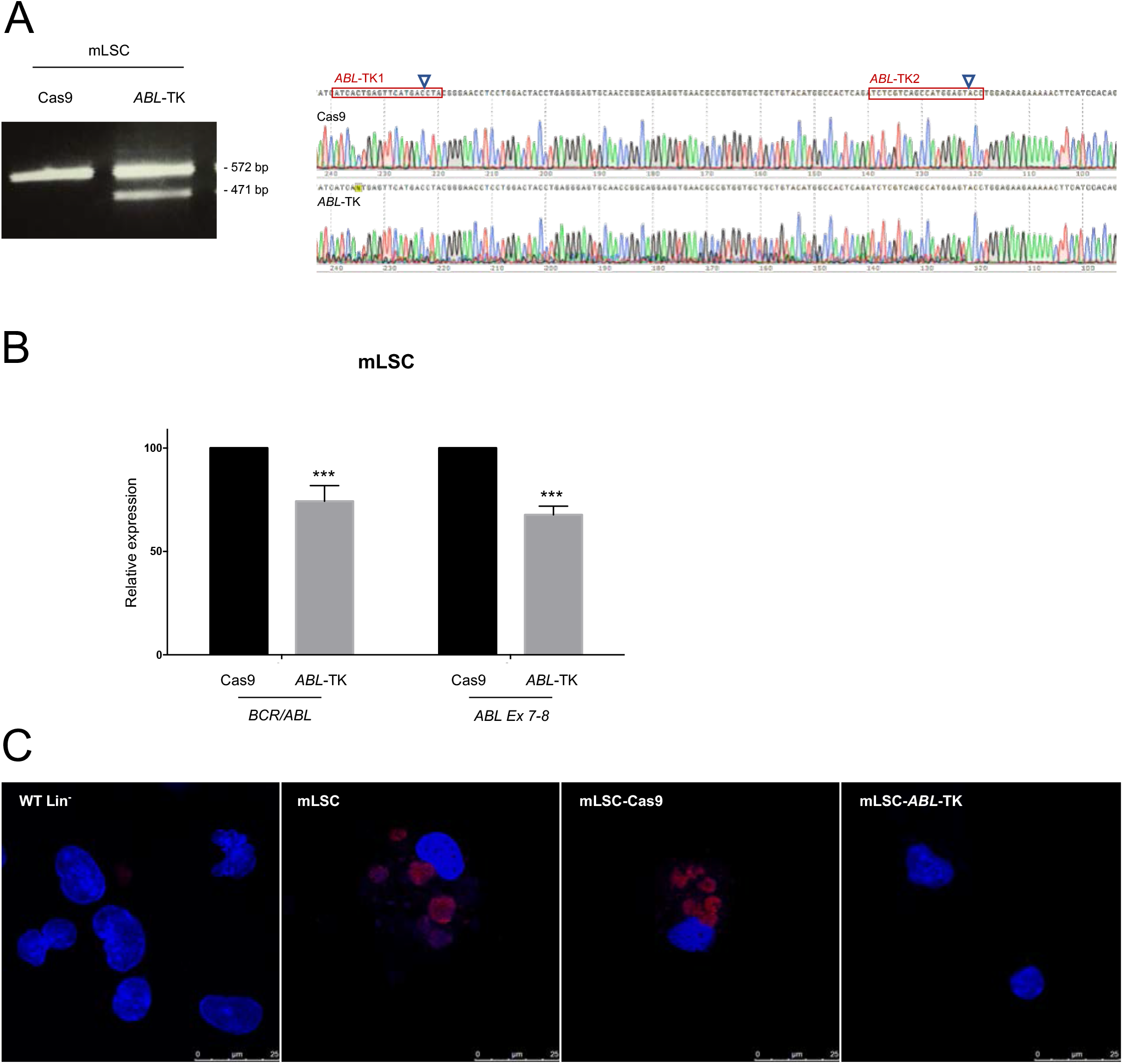
CRISPR/Cas9 deletion system in mouse leukemic stem cells. Mouse leukemic stem cells (mLSCs) were isolated from TgP210+ mouse bone marrow, and electroporated with Cas9 and with Cas9 joined to both sgRNAs (*ABL*-TK). (A) PCR-amplification of the genomic ABL exon 6 sequence and Sanger sequencing. (B) Quantitative PCR of *BCR/ABL* mRNA (fusion point) and *ABL* mRNA (exon 7-8) in mLSC-Cas9 (control) and mLSC-*ABL*-TK (edited) cells (mean ± SEM; ***p<0.001). (C) Anti-human ABL immunostaining of mLSC mouse cells.

Total mRNA from electroporated mLSCs-*ABL*-TK cells was isolated and the expression level of BCR/ABL quantified by qPCR (Figure 3B). While mLSCs-Cas9 control cells showed high BCR/ABL mRNA levels, we found significantly (p<0.001) lower levels of expression in mLSCs-Cas9 cells. Accordingly, a markedly lower levels of ABL protein were detected by immunofluorescence in mLSCs-*ABL*-TK cells (Figure 3C).

To test the multipotency of these mLSCs-*ABL*-TK cells we transplanted them and their counterpart mLSCs-Cas9 cells into two irradiated immunodeficient NSG mice (NSG-Cas9 and NSG-*ABL*-TK; Figure 4A). Several hematopoietic cell lineages from the peripheral blood of transplanted NSGs were analyzed 120 days post-transplant (Figure 4B). Significantly, we detected and isolated specific hematopoietic cells populations such as myeloid cells (Gr1^+^), B lymphocytes (B220^+^) and T lymphocytes (CD4^+^ or CD8^+^), arising from both NSG-*ABL*-TK and NSG-Cas9 mice. In contrast, FACs analysis only detected normal white cell percentages in peripheral blood from NSG-*ABL*-TK, while the NSG-Cas9 counterpart exhibited a Gr1^+^ bias to the detriment of the lymphoid linage (Figure 4B). As expected, PCR amplification of *ABL* exon 6 in all sorted populations from NSG-*ABL*-TK showed the presence only of the CRISPR/Cas9 *ABL1* (Figure 4C). Importantly, compared with NSG-Cas9, a smaller Gr1^+^ population was detected in NSG-*ABL*-TK, at least during the first 60 days post-transplant. The *BCR/ABL* qPCR assay in sorted CD45^+^ cells from NSG-*ABL*-TKs at 120 days post-transplant showed a lower level of expression (69.1 ± 1.9%) than its NSG-Cas9 counterpart (Figure 4D).

**Figure 4.**
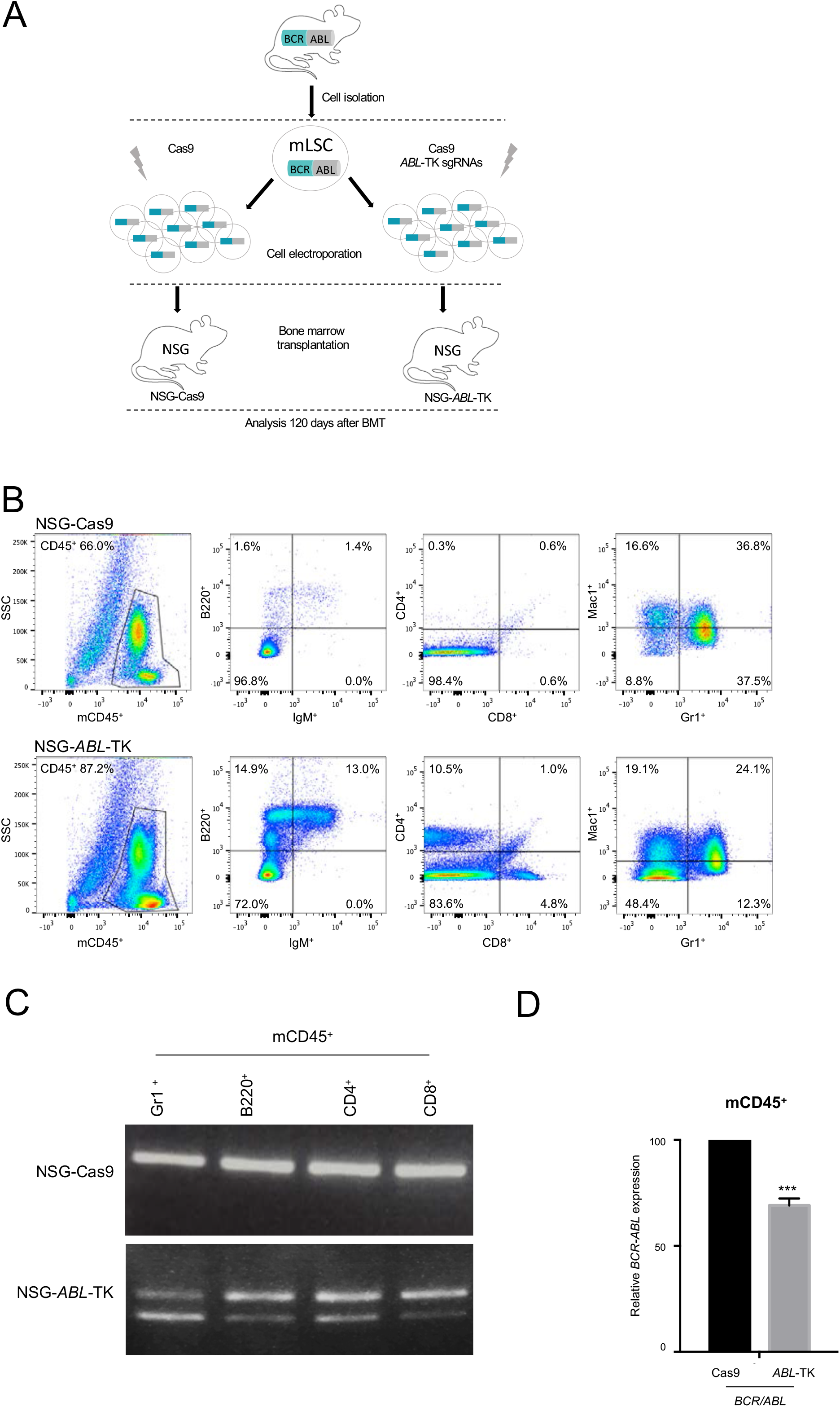
Engraftment and Multipotency capacity of CRISPR/Cas9 edited mLSCs. (A) Schematic representation of the experimental procedure and study groups. (B) 4 months after BMT, mouse CD45^+^ cell populations from peripheral blood were analyzed and isolated by FACS. Lymphoid (B220^+^, CD4^+^, and CD8^+^) and myeloid (Gr1^+^ and Mac1^+^) populations were identified in the blood samples. (C) DNA samples from mCD45^+^ cells from mLSC-Cas9 and mLSC-*ABL-*TK mice were PCR-amplified, revealing the presence of a CRIPSPR/Cas9-induced deletion in *ABL* exon 6 of the GR1^+^, B220^+^, CD4^+^, CD8^+^ and SCA^+^ cells. (D) Quantification of *BCR/ABL* expression in recovered mCD45 cells 4 months after transplantation (mean ± SEM; ***p<0.001).

### 4. mLSCs modified by CRISPR at the ABL1 locus impairs the myeloid bias, restoring normal hematopoiesis

To determine whether LSC-*ABL*-TK could be potentially useful as a CML therapy, 12 independent NSG bone marrow transplantation assays of mLSCs CRISPR-edited and non-edited were performed (Figure 5). Four groups of mice were studied: 1) six NSG mice with CRISPR-edited mLSCs from CML donors (NSG-*ABL*-TK), 2) six NSG mice with non-edited mLSCs from CML donors (NSG-Cas9), 3) five CML transgenic mice, used as parental disease controls (CML-Tg), and 4) five wildtype mice, used as normal controls (WT) (Figure 5A). All mLSCs were isolated from bone marrows of at least one-year-old CML transgenic mice with clinical disease (>60% Gr1^+^ cells in peripheral blood) (Supplementary table 2). The *BCR/ABL* transgene sequence was CRISPR-disrupted in Lin^−^ mLSCs, as described above. CRISPR/Cas9-specific deletion at the *ABL1* TK domain was confirmed by PCR and Sanger sequencing. Peripheral blood cell analysis by FACs was performed 60 days after transplant to study the hematological cell populations. The end-point was determined at 120 days (Supplementary table 3).

**Figure 5.**
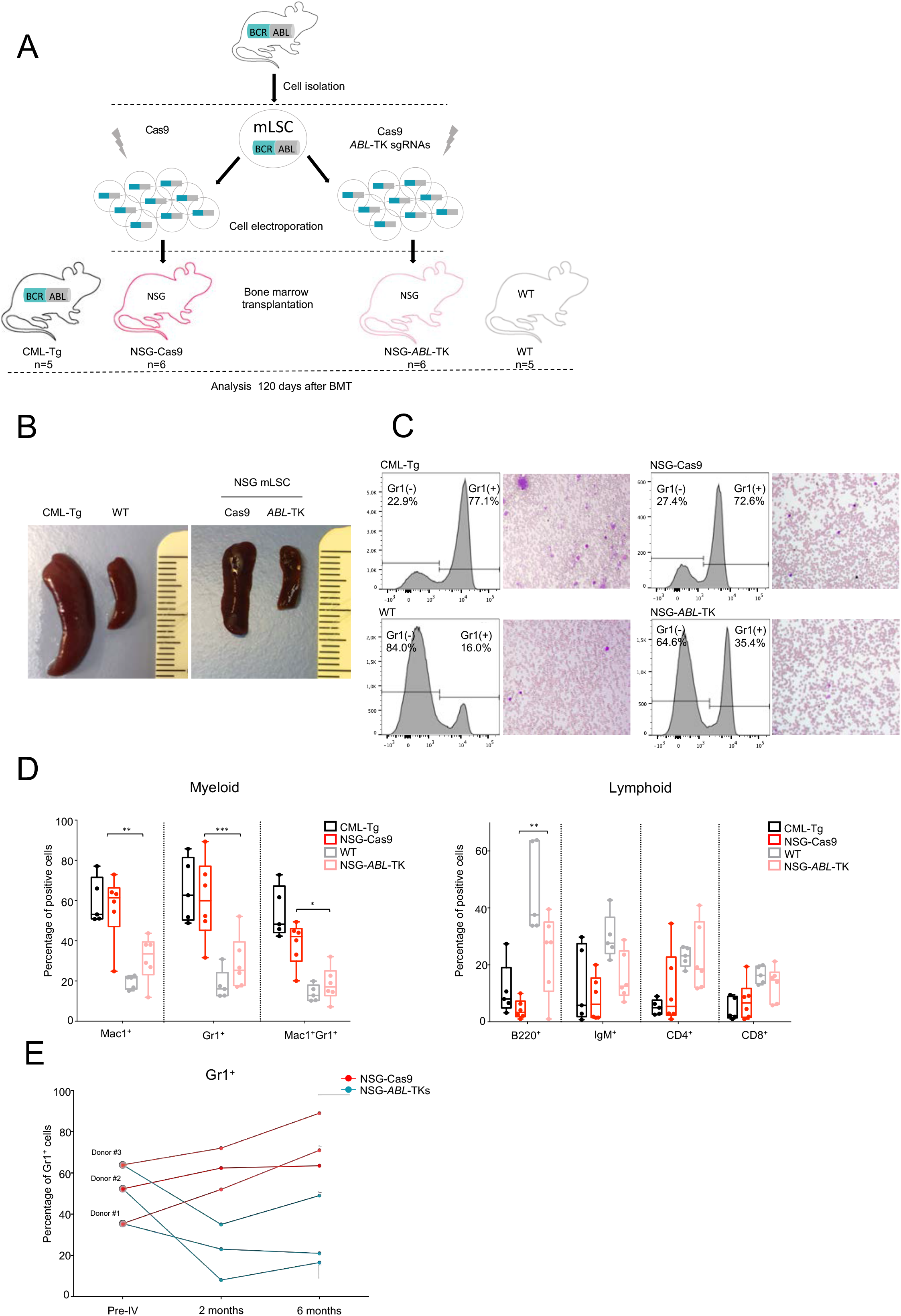
Therapeutic evaluation of CRISPR/Cas9 deletion system in a Mouse model of LMC. (A) Schematic representation of the experimental procedure and study groups. (B) Macroscopic aspect and spleen sizes of CML transgenic mice, wild-type mice, and NSG recipient mice (NSG-Cas9 and NSG-*ABL*-TK). (C) FACS quantification of Gr1-positive cells and representative micrography of hematoxylin-stained peripheral blood. **(D)** FACS analysis of myeloid and lymphoid hematological populations generated in peripheral blood of NSG recipient mice after BMT: NSG-*ABL*-TK; (pink box), NSG-Cas9 (red box). CML transgenic mice (black box) and wild-type mice (grey box) were used as controls for CML disease and normal cell distribution (median ± SEM; *p<0.05; **p<0.01; ***p<0.001). **(E)** Percentage and evolution of Gr1^+^ cell population analyzed in peripheral blood of NSG recipients with mLSC engraftments from CML transgenic mice at different stages disease. mLSCs were divided into two groups with the same number and were electroporated with Cas9 (NSG-Cas9, red line) and with Cas9 joined to both sgRNAs (NSG-*ABL*-TK, blue line) before BMT.

We observed that four of the six NSG-Cas9 mice suffered splenomegaly, as was observed in CML transgenic mice. However, the spleens of all the NSG-*ABL*-TK were of normal size compared with those of WT mice (Figure 5B).

The proportion of Gr1^+^ cells increased in all NSG-Cas9 mice, reaching 59.2 ± 9.9% in peripheral blood by the end-point. In contrast, a significantly lower percentage of Gr1^+^ cells was detected in NSG-*ABL*-TKs (29.2 ± 6.6; p<0.001), similar to what was observed in WT mice (17.9 ± 3.4) (Figure 5C-5D). There was a significantly higher percentage of Mac1^+^ cells (55.2 ± 8.2) in NSG-Cas9 than the normal levels seen in NSG-*ABL*-TK (31.7 ± 5.7; p<0.05) and WT mice (19.6 ± 1.5; p<0.05). As expected, there were low levels of aberrant lymphoid cell populations in NSG-Cas9 and CML mice (Figure 5C-D).

To study in greater detail the how the time to evolution of the Gr1^+^ cell population is related to disease stages while accounting for interindividual differences among donors, three independent transplantation assays were performed with three independent mLSC donor mice (Supplementary table 2; Figure 5E). The different clinical disease stages were defined as “initial phase” (donor #1, Gr1^+^ ≤40%), “chronic phase” (donor #2, Gr1^+^ ≥40% and ≤60%), and “advanced phase” (donor #3, Gr1^+^ ≥ 60%). LSCs from each CML mouse were divided in two samples and electroporated with or without all CRISPR reagents (NSG-*ABL*-TK and NSG-Cas9). Both samples were transplanted into NSG receptors and maintained for 6 months (Figure 5E). At the time of transplantation, the CML donor mice selected showed 35.4% (donor#1), 52.3% (donor#2) and 63.9% (donor#3) of Gr1^+^ cells. 4 months after transplantation, all NSG-*ABL*-TKs mice had normal levels of Gr1^+^ cells (donor#1, 23%; donor#2, 8%; donor#3, 35%) and were similar 6 months after transplantation. In contrast, all NSG-Cas9 mice showed increasing Gr1^+^ populations at 2 months and reached pathological levels by the end-point (donor#1, 52%; donor#2, 62%; donor#3, 72%) (Figure 5E).

### 5. The CRISPR/Cas9 deletion system efficiently disrupts the BCR/ABL oncogene in CD34^+^ LSCs from CML patients, impairing the myeloid bias and restoring its physiological multipotency

To corroborate the applicability of CRISPR technology as a potential therapeutic tool in human CML, LSCs CD34^+^ were isolated from CML patients at diagnosis in the chronic phase. Similar to the assays described above, we electroporated CD34^+^ hLSCs with Cas9 RNP, with or without two specific sgRNAs targeting the *BCR/ABL* exon 6 (hLSCs-*ABL*-TK and hLSCs-Cas9). A 473-bp band, corresponding to *BCR/ABL* genomic exon 6, and an extra band of 372 bp were also detected by PCR in hLSC-*ABL*-TKs. As expected, a single 473-bp band was amplified in hLSCs-Cas9 and control CD34^+^ cells (Figure 6A). Sanger sequencing confirmed the specific deletion between the expected Cas9 cut sites (Figure 6A). Quantitative PCR of mRNA from hLSCs-*ABL*-TKs showed a significantly lower level of expression of BCR/ABL mRNA (p<0.001) than in controls (Figure 6B). These results were confirmed by western blot (Figure 6C).

**Figure 6.**
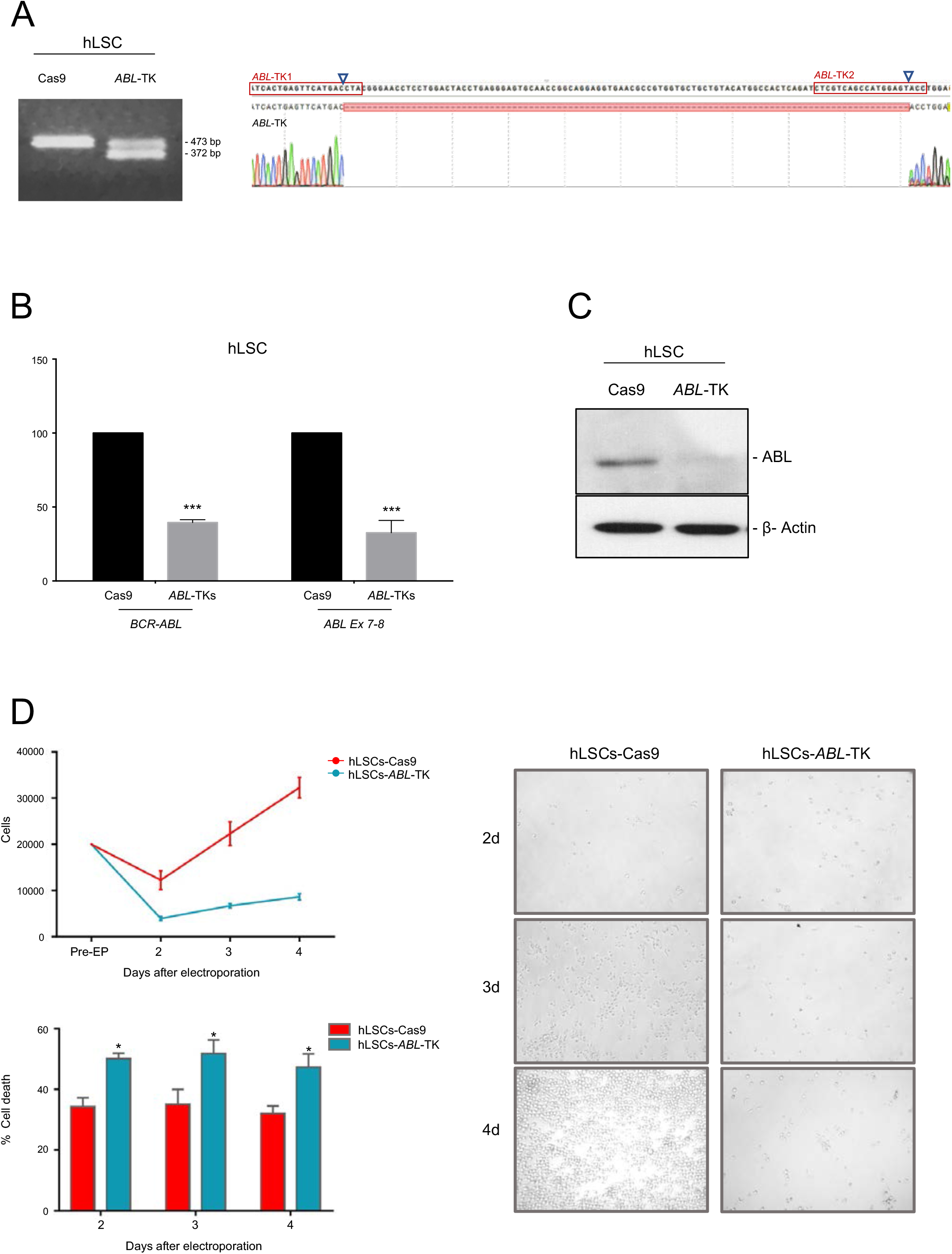
Effects of CRISPR/Cas9 deletion system in human LSCs. Human CD34^+^ LSCs were isolated from bone marrow biopsies of CML patients, and electroporated with/without the all CRISPR/Cas9 reagents. **(A)** PCR amplification of ABL exon 6 in hLSC from patients electroporated with Cas9 nuclease (hLSC-Cas9) and Cas9 joined to both sgRNAs (hLSC-*ABL*-TK). Sanger sequencing of ABL exon 6 of both cell populations. **(B)** Quantitative PCR of *BCR/ABL* (fusion *BCR/ABL* exons, left) and *ABL* (exon 7-8, right) (mean ± SEM; *** p<0.001). **(C)** Western blot anti-ABL in hLSC cells. Cells electroporated with Cas9, used as control, showed a single band corresponding to ABL (~135 kDa). However, hLSC electroporated with *ABL*-TKs showed a lower level expression level of ABL protein. **(D)** *In vitro* cell proliferation and apoptosis analysis of hLSC after CRISPR/Cas9 editing. LSC proliferation and apoptosis levels were measured 24, 48 and 96 h after electroporation with Cas9 (red) and Cas9 joined to ABL-TK sgRNAs (blue)(mean ± SEM;*p<0.05). hLSC-*ABL*-TK showed a lower proliferation ability and a higher apoptosis rate than non-edited cells (hLSC-Cas9).

To explore the ability of the CRISPR/Cas9 system to abolish the tumorigenic capacity of hLSCs, we studied proliferation and survival of hLSC-*ABL*-TK cells *in vitro* (Figure 6D). hLSC-*ABL*-TK had a lower proliferation rate than did hLSC-Cas9 cells. Accordingly, hLSC-*ABL*-TK also showed a higher percentage of cell death than did hLSC-Cas9 cells (Figure 6D).

Similar to NSG bone marrow transplantation assays described above, CML CD34^+^ cells from 10 patients at diagnosis in CP-CML (7 of them with enough CD34^+^ cells to split in 2 samples each one) were included to establish orthotopic PDX assays in NSG mice as follow: 7 CML CD34^+^ samples that were electroporated with Cas9 and *ABL1* sgRNAs (hLSC-*ABL*-TKs), 7 CML CD34^+^ that were electroporated solely with Cas9 (hLSCs-Cas9), 3 un-electroporated CML CD34^+^ cells as disease controls. 3 CD34^+^ samples from healthy donors were used as controls (NSG-healthy-hCD34^+^). A total of 20 NSG mice of the same age were used as receptors of these cells (Figure 7A). A representative FACS analysis of bone marrow 180 days after transplantation is shown in Figure 7B. In all cases, the percentages of myeloid cells (hCD14 and hCD117) and lymphoid B cells (hCD19) in NSG-hLSC-*ABL*-TK were normal. mRNA from hCD45^+^-sorted bone marrow cells was isolated from NSG-hLSC-Cas9 and NSG-ABL-TKs mice to analyze *BCR/ABL* expression by qPCR (Figure 7C). NSG-hLSC-*ABL*-TK presented significantly lower levels of expression of BCR/ABL and ABL (30.3 ± 12.5% and 33.2 ± 17.7%, respectively; p<0.01) than did NSG-hLSC-Cas9 mice (Figure 7C).

**Figure 7.**
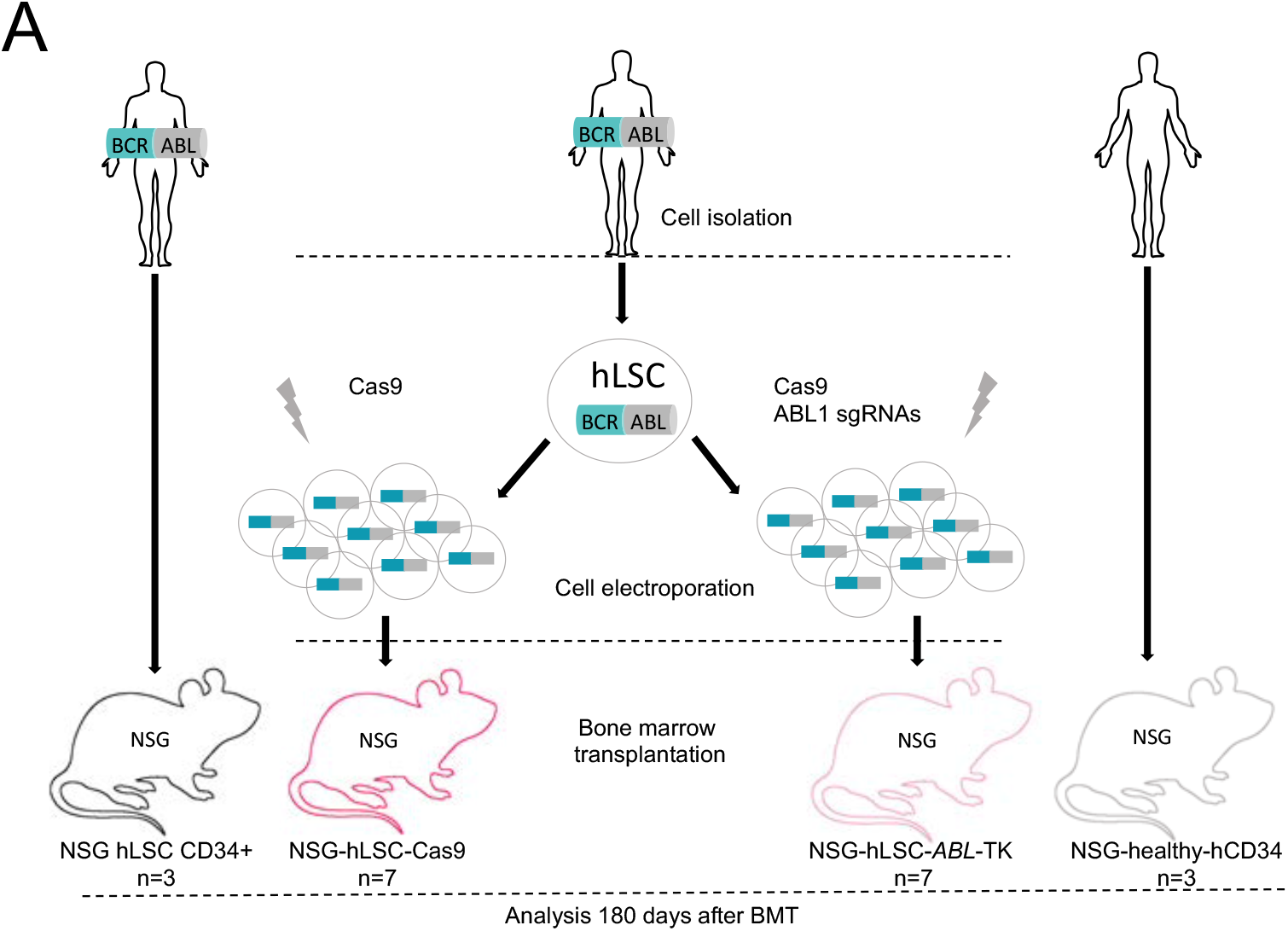
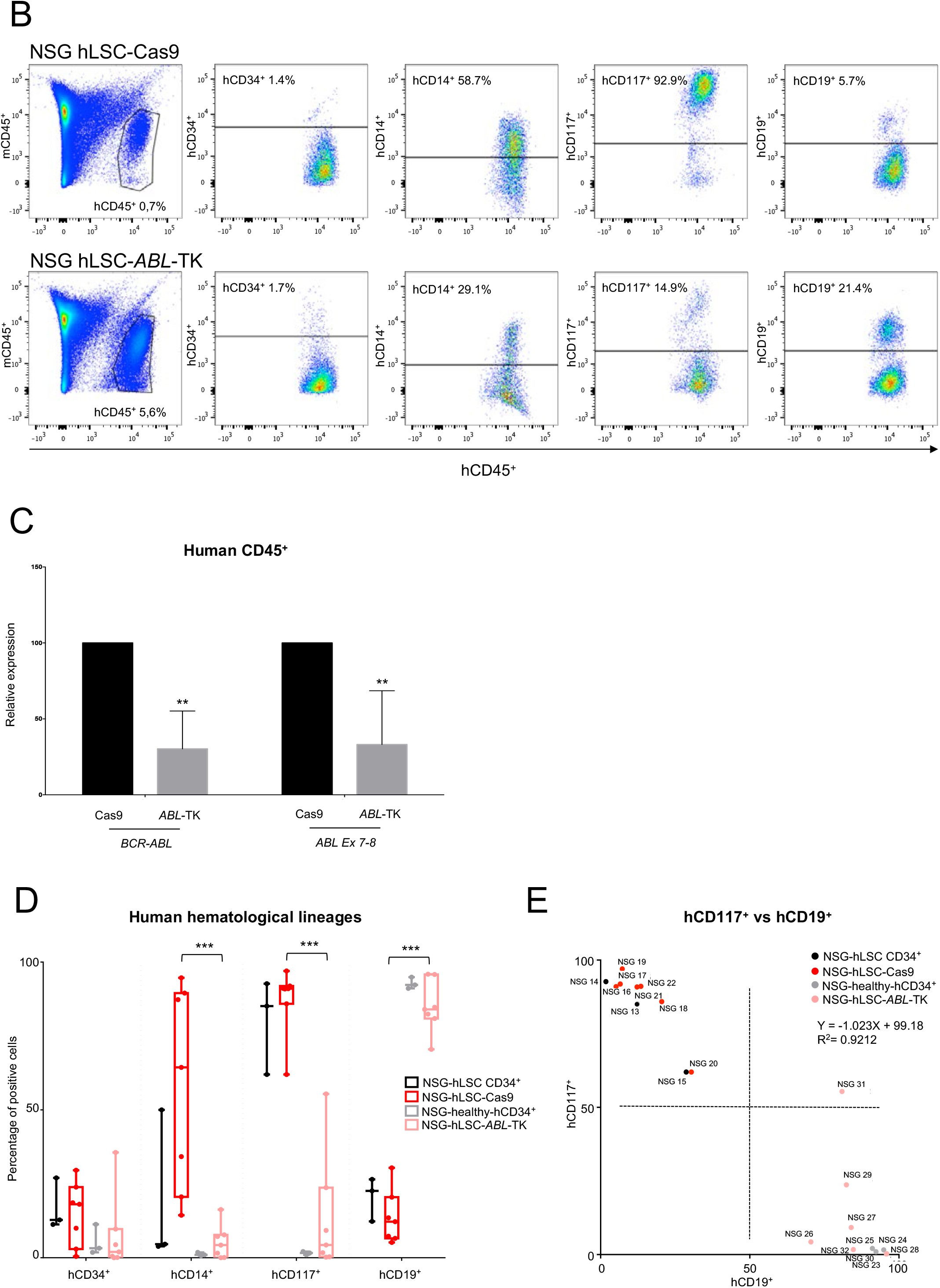
Multipotency capacity and therapeutic evaluation of CRISPR-edited hLSCs in an orthotopic model. Human CD34^+^ LSCs were isolated from bone marrow biopsies of CML patients, electroporated with/without all CRISPR/Cas9 reagents and transplanted into irradiated NSG mice. **(A)** Schematic representation of study groups and experimental procedure. **(B)** FACS analysis of human hematological cell populations in bone marrow 24 weeks after BMT. NSG with hLSC engraftments from patients electroporated with Cas9 nuclease are named NSG-hLSC-Cas9. NSG with hLSC engraftments from patients electroporated with Cas9 joined both sgRNAs are named NSG-hLSC-*ABL*-TK. **(C)** Quantitative PCR of *BCR/ABL* (fusion *BCR/ABL* exons, left) and *ABL* (exon 7-8, right) in hCD45 cells from NSG bone marrow 24 weeks after BMT. **(D)** FACS analysis of human hematological hCD45^+^ cell population 24 weeks after BMT in NSG mice. NSG with hLSCs engraftments electroporated with Cas 9 are named NSG-hLSC-*Cas9* (red boxes) and with Cas9 joined to sgRNAs are named hLSC-*ABL*-TK (pink boxes). NSG with healthy hCD34^+^ engraftments (NSG-healthy-hCD34^+^, grey boxes) were used as normal controls. NSG with untreated hCD34^+^ cells from CML patient (NSG-hLSC-CD34^+^; black boxes) were used as a control disease. (***p<0.001). **(E)** Linear regression analysis of hCD117+ and hCD19+ cell populations in the NSG mice engraftments. NSG with hLSCs engraftments electroporated with Cas 9 are represented with pink dots, with Cas9 joined to sgRNAs are represented with red dots. NSG with healthy hCD34^+^ engraftments (NSG-healthy-hCD34^+)^ are represented with grey boxes. NSG with untreated hCD34^+^ cells from CML patient (NSG-hLSC-CD34^+^) are represented with black boxes.

FACS analysis of human hematological populations from NSG bone marrow samples was carried out 180 days after transplantation (Figure 7D, Supplementary table 4), which revealed normal levels of hCD14^+^ (5.4 ± 2.6%), hCD117^+^ (15.5 ± 8.7%) and hCD19^+^ (84.9 ± 3.9%) in all NSG-hLSC-*ABL*-TK. In contrast, higher levels of CD14^+^ (65.1 ± 12.7%) and CD117^+^ (86.5 ± 5.1%), and reduced levels of hCD19^+^ (15.1 ± 3.7%) were detected in all NSG-hLSC-Cas9 bone marrow samples (p<0.001) (Figure 7D).

Importantly, a linear regression analysis of hCD117^+^ and hCD19^+^ cell populations in the NSG mice engraftments showed an inverse correlation (r^2^ = 0.9212) among NSG-hLSC-*ABL*-TK and NSG-healthy-hCD34+ with respect to NSG-hLSC-Cas9 and NSG-hLSC CD34^+^ donors (Figure 7E).

## DISCUSSION

As our understanding of the molecular pathogenesis of CML has developed, its prognosis has improved, from being that of a fatal disease, to one of a chronic disorder that can be treated with oral medication that could be even discontinued in a small subset of patients ^48^. This change has been possible due to the development of targeted drugs, such as TKIs, that inhibit the oncoprotein ^23^. Unfortunately, these drugs do not tackle the fundamental cause of the disease, and the oncogenic event remains unaffected or uncorrected. Thus, lifelong oral medication is necessary for most CML patients, and a significant percentage of patients eventually become resistant to TKI treatment ^49^.

Currently, an oncogenic event can be easily abolished by using CRISPR/Cas9 nucleases, which offers a new and definitive opportunity for TKI-resistant CML patients ^32,33^. Our previous CRISPR studies and later others studies with CRISPR RNA-guided engineered with FokI nucleases ^35^ or CRISPR/Cas9 nucleases delivered by lentivirus ^37^, demonstrate the effectiveness of CRISPR/Cas9 system for knocking out *ABL1*, the key oncogene in the CML disease^34^, avoiding its oncogenic potential similarly to described with TKI imatinib^50^. All these previous studies have in common the induction of insertion/deletions (indels) at *ABL1* gene to interrupt the coding sequence for abolishing its expression in a permanent manner. Shu-Huey Chen et al.^37^ isolated transduced peripheral blood mononuclear cells (PBMCs) of a CML patient with an ABL1 sgRNA-CRISPR/Cas9 lentivirus. They demonstrated a significant increase in the number of cells undergoing apoptosis after transduction. We corroborated their findings, for the first time, in *ABL1* CRISPR-edited primary bone marrow Lyn^−^LSCs isolated from CML transgenic mice and *ABL1* CRISPR-edited primary CD34^+^LSCs from CP-CML patients. The absence or reduced level of BCR/ABL1 expression was corroborated at the protein level and triggered cellular apoptosis, particularly in the edited CML cell lines, and arrested proliferation, especially in primary edited LSC samples. Recently, a similar approach with zinc finger nucleases have been tested but to disrupt *BCR* exon1 and adding 8-base NotI enzyme cutting site by homology-directed repair, which led to a stop codon and truncated BCR-ABL protein^36^. Unlike all these papers and based on previous studies of Bauer DE et al., we have developed a new strategy to generate CRISPR induced deletions at *ABL1* locus. Deletions have potential advantages as compared to single-site small indels given the efficiency of biallelic modification, ease of rapid identification by PCR, predictability of loss-of-function^51^. To ensure a null oncogenic effect, here we have designed two sgRNAs to induce a specific short deletion at the TK domain of *ABL1*, which guarantees the production of a truncated protein with no TK activity. Furthermore, this deletion allows the tracking of edited cells and their daughter cells. Our results indicate that CRISPR/Cas9 system with a dual guide could be used to generate ABL1 null alleles and to abolish BCR/ABL1 transcripts preventing any oncogenic effect on survival and proliferation specially when edited cell can be selected or isolated to grow. It is important to remark that a remaining expression of BCR/ABL is observed in all studies with primary mouse or human LSCs, in which it is not possible to select correctly edited cells, and the residual expression is due to the overall efficiency ratio intrinsic to the delivery and genome edition processes^32,33,35–37^. Despite this, a huge reduction of BCR/ABL1 transcripts was achieved with a significantly therapeutic benefit in mouse models and PDX.

In other side, critical questions about using CRISPR as a new potential tool for CML treatment remain, such as whether CRISPR-edited LSCs can engraft and dominate bone marrow hematopoiesis or, if the edited LSC recovers its physiological multipotential commitment and thereby avoids the myeloid bias. Mandal et al showed that a dual guide approach improved gene deletion efficacy in healthy CD34+ hematopoietic stem and progenitor cells preserving the multilineage potential of the CD34^+^ cell^52^. In primary human CD4^+^ T cells and CD34^+^ hematopoietic stem and progenitor cells, they demonstrated that multi-lineage potential of the CD34^+^ cell is retained when the cell is CRISPR-edited. Nevertheless, in a CRISPR-edited leukemic stem cell these are unresolved questions yet. To address these questions, we performed several mouse-mouse and human-mouse BMT assays using immunodeficient NSG mice as the best recipient for studying the multipotency of human and mouse hematopoietic stem cells *in vivo*^53^. A humanized transgenic mouse for mimicking CML was used as an LSC donor to test the engraftment capacity and multipotency of CRISPR-edited mLSCs. In contrast to NSG-Cas9 controls, we detected physiological percentages of myeloid and lymphoid cells in NSG-*ABL*-TK due to the multilineage capacity of the CRISPR-edited mLSC. Importantly, myeloid bias was not detected in NSG-*ABL*-TK, at least 120 days after BMT, which suggests that CRISPR system with dual guide significantly reduces the tumor burden, given rise to hematological populations with physiological percentages. No clinical symptoms of splenomegaly and increased granulocyte lineage, that are usually reported when mice receiving P210 bcr/abl-transduced bone marrow cells^54–56^ were detected in all NSG-*ABL*-TKs. In contrast, we observed identical symptoms in NSG-Cas9 and CML-Tg mice. To study the evolution of BMTs with respect to disease stage, which is closely related to tumor burden, we measured the Gr1^+^ percentage at the beginning, and 60 and 180 days post-BMT. We detected higher percentages of Gr1^+^ tumoral cells in all NSG-Cas9 controls, suggesting a CML development. In contrast, 180 days post-BMT, all NSG-*ABL*-TK mice showed lower levels of Gr1^+^ cells than those detected at the start. Therefore, we concluded that the CRISPR deletion system restored normal hematopoiesis and produced a therapeutic benefit in the CML mouse model. To determine whether that result was reproducible, similar BMT assays were performed with CML human samples collected from 10 untreated CP-CML patients. Based on previous papers, successful engraftment was defined as the presence of at least 0.1% human CD45^+^ cells in bone marrow, measured by flow cytometry^57^. All NSG mice included in this study presented engraftments between 0.7 % and 9.4 %, whereby migration and self-renewal capacity of the transplanted cells (edited and non-edited) were not affected. Like Mandal et al.^52^, we noted the presence 120 days after BMT of CD45^+^ cells arising from the edited mLSCs. PCR analysis on DNA isolated from sorted human CD45^+^ hematopoietic cells from reconstituted mice demonstrated that *ABL1* edited cells robustly contributed to human hematopoietic cell chimerism. As expected, we detected normal hematopoiesis in all NSG-hLSC-*ABL*-TK mice, analogous to what was found in NSG-healthy CD34^+^ cases. Similar to other authors have noted in NSG with human HSC engraftments from healthy controls ^58^, we also observed a normal hematopoiesis in all NSG-hLSC-*ABL*-TK mice defined by a high proportion of B cells, very low levels of T cells, and the absence of myeloid bias. In contrast, all NSG-hLSC-Cas9 developed a myeloproliferative disorder in a similar way as occurs in NSG-hLSC CD34^+^, with a high proportion of CD14^+^ and CD117^+^ cells. CD117 and CD19 expression were negatively correlated when all the engraftments at 180 days were analyzed. High CD117 and low CD19 populations were found in NSG-hLSC-Cas9 mice, while an inverse result was observed in NSG-hLSC-ABL-TK. Since a close inverse correlation between CD117 and CD19 expression was associated with the myeloid phenotype and/or the progression of CML by W Eisterer et al.^59^, our results indicate that the CRISPR/Cas9 deletion system is able to abolish the oncological effect of BCR/ABL expression, given rise a corrected hematological differentiation after BMT. More importantly, the CRISPR-edited LSC cells retain the cells the ability to restore normal hematopoiesis avoiding the myeloid bias. The treatment has been successfully evaluated in mouse models and in PDX. In our opinion, CML could be an ideal candidate for CRISPR therapy in those cases for which current TKI line of therapies are not effective.

Taken together, our results constitute the first step towards providing proof-of-principle for genome editing of the main genetic event in CML patients, and suggest that CRISPR/Cas9 technology could provide the basis for a very useful alternative drug for treating human CML.

## ACKNOWLEDGEMENTS

We thank the Radioactive Isotopes and Radioprotection Services for mouse irradiation, and the cell separation, experimental animal and transgenic facilities of NUCLEUS, University of Salamanca, for carrying out the FACS and animal assays.

## FUNDING

This work was supported by the Research Support Platform of the University of Salamanca (NUCLEUS), Instituto Carlos III (ISCIII) PI17/01895 (ISCIII-FEDER), Novartis Farmaceutica S.A., a predoctoral grant from University of Salamanca-Banco Santander, and Jabones Solidarios para Daniel from the Bomberos Ayudan Association.

## AUTHOR CONTRIBUTIONS

**Conceptualization**: Manuel Sánchez-Martín.

**Design and Investigation**: Ignacio García-Tuñón, Manuel Sánchez-Martín.

**Data curation**: Elena Vuelta, Ignacio García-Tuñón.

**Statistical analysis**: Elena Vuelta, Ignacio García-Tuñón.

**Methodology:** Elena Vuelta, Ignacio García-Tuñón, Manuel Sánchez-Martín.

**Animal care**: Lucía Méndez, Patricia Hernández-Carabias, José Luis Ordóñez.

**Resources:** Lucía Méndez, Patricia Hernández-Carabias, José Luis Ordoñez, Verónica Alonso-Pérez.

**Human samples and clinical data:** Raquel Saldaña, Julián Sevilla, Elena Sebastian, Fermín Sánchez-Guijo, Sandra Muntión, Jesús María Hernández-Rivas.

**Funding acquisition**: Jesús María Hernández-Rivas, Ignacio García-Tuñón, Manuel Sánchez-Martín.

**Project administration:** Ignacio García-Tuñón, Manuel Sánchez-Martín.

**Supervision:** Ignacio García-Tuñón, Manuel Sánchez-Martín

**Figures:** Elena Vuelta, Ignacio García-Tuñón, Manuel Sánchez-Martín

**Writing – original draft:** Ignacio García-Tuñón, Jesus María Hernández-Rivas, Manuel Sánchez-Martín.

## Competing interests

The authors have declared that they have no competing interests.

---

**Supplementary Figure 1.**
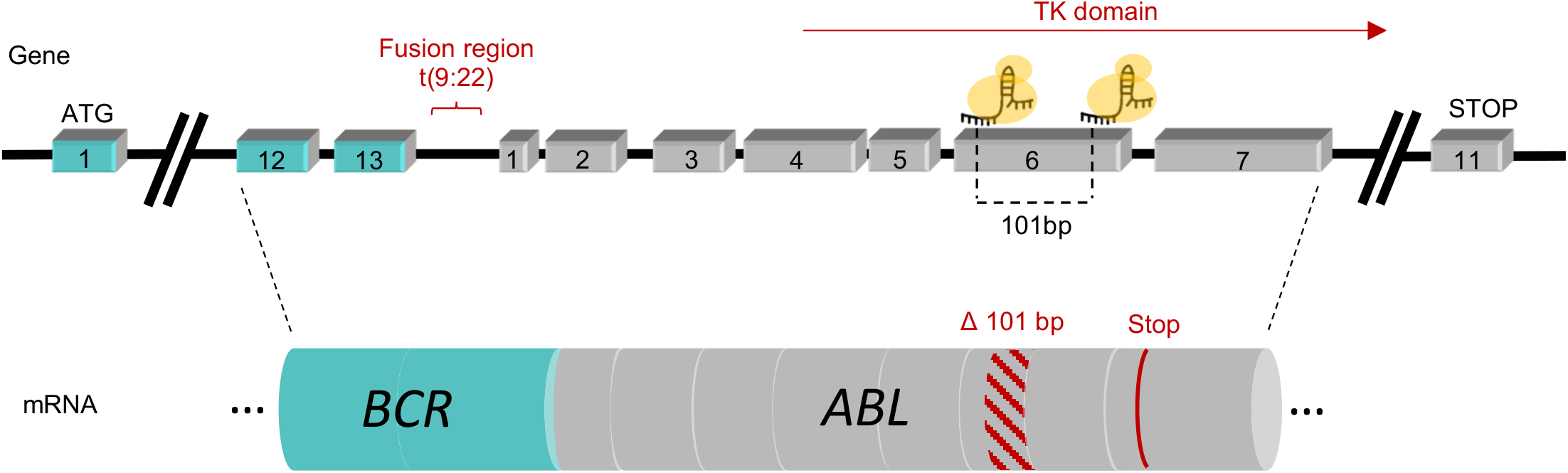
Experimental design of the CRISPR/Cas9 system for blocking the *BCR/ABL* fusion oncogene. Detail of fusion *BCR/ABL* gene resulting from t(9:22) and the expected cleavage point of sgRNA against *ABL* exon 6. The resulting 101-bp deletion would generate a premature stop codon in exon 7.

**Supplementary Figure 2.**
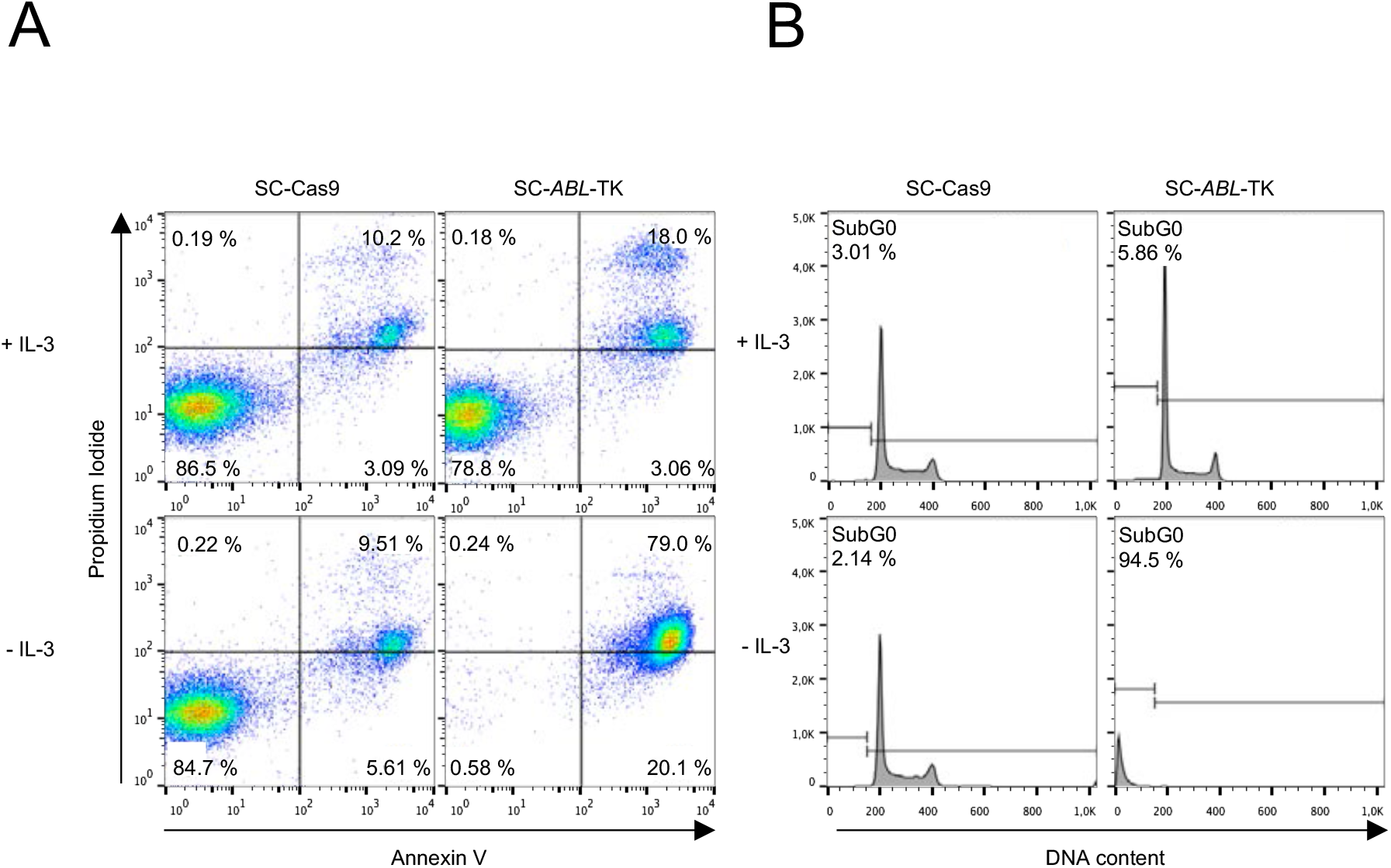
Functional analysis of CRISPR/Cas9 deletion system in a CML murine cell line. (A) Annexin-V cell apoptosis assay of SC cells in the presence or absence of IL-3. (B) Cell cycle assay of SC cells in the presence or absence of IL-3.

**Supplementary Figure 3.**
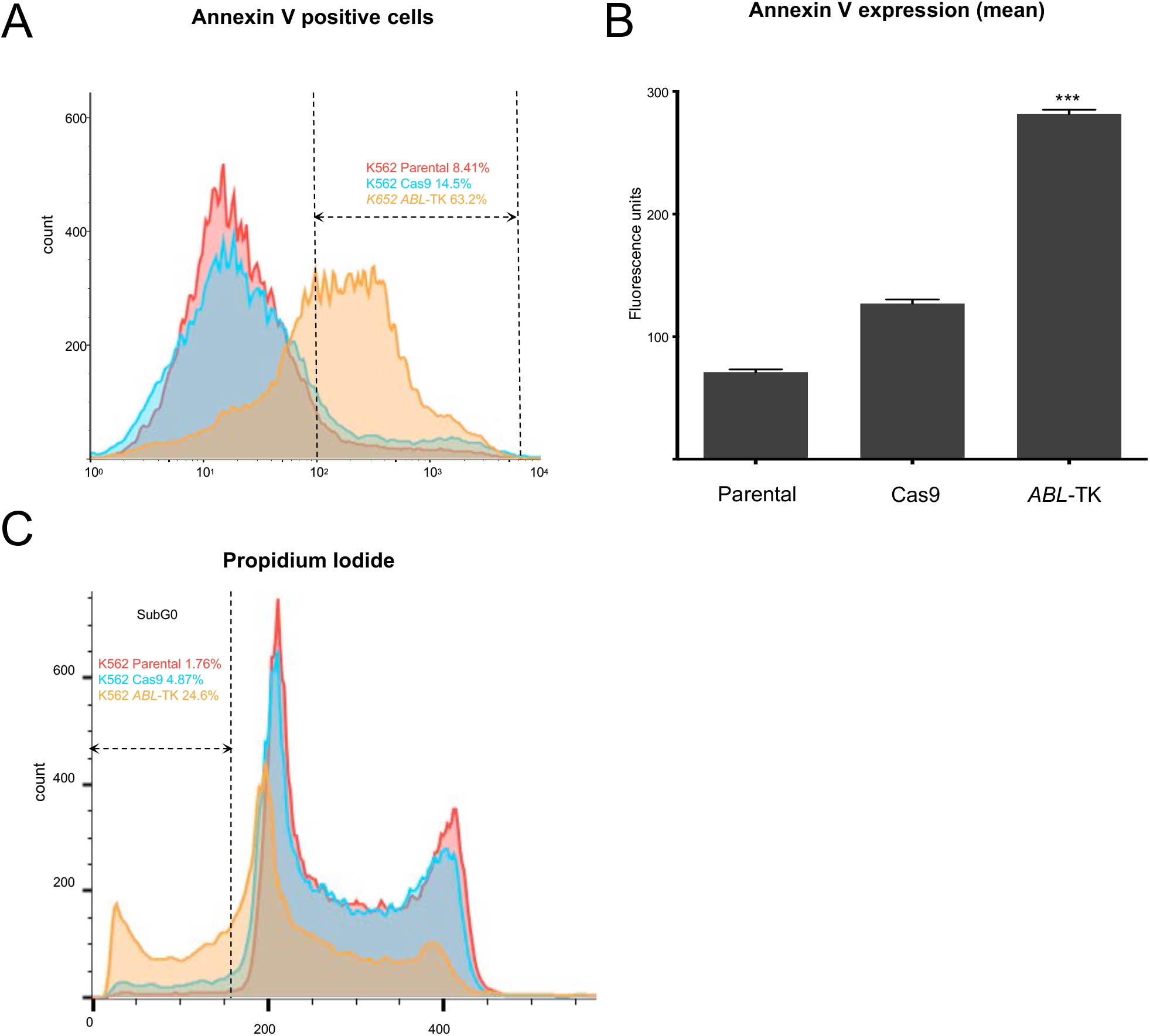
Analysis of cell viability in the CRISPR/Cas9-deletion system in a CML human cell line. (A) Annexin-V cell apoptosis assay of parental, K562-Cas9 and – K562-*ABL*-TK cells. (B) Quantification of annexin-V labeling in cell populations (mean ± SEM; *** p<0.001). (C) DNA content analysis of cell populations.

**Supplementary Table 1.**
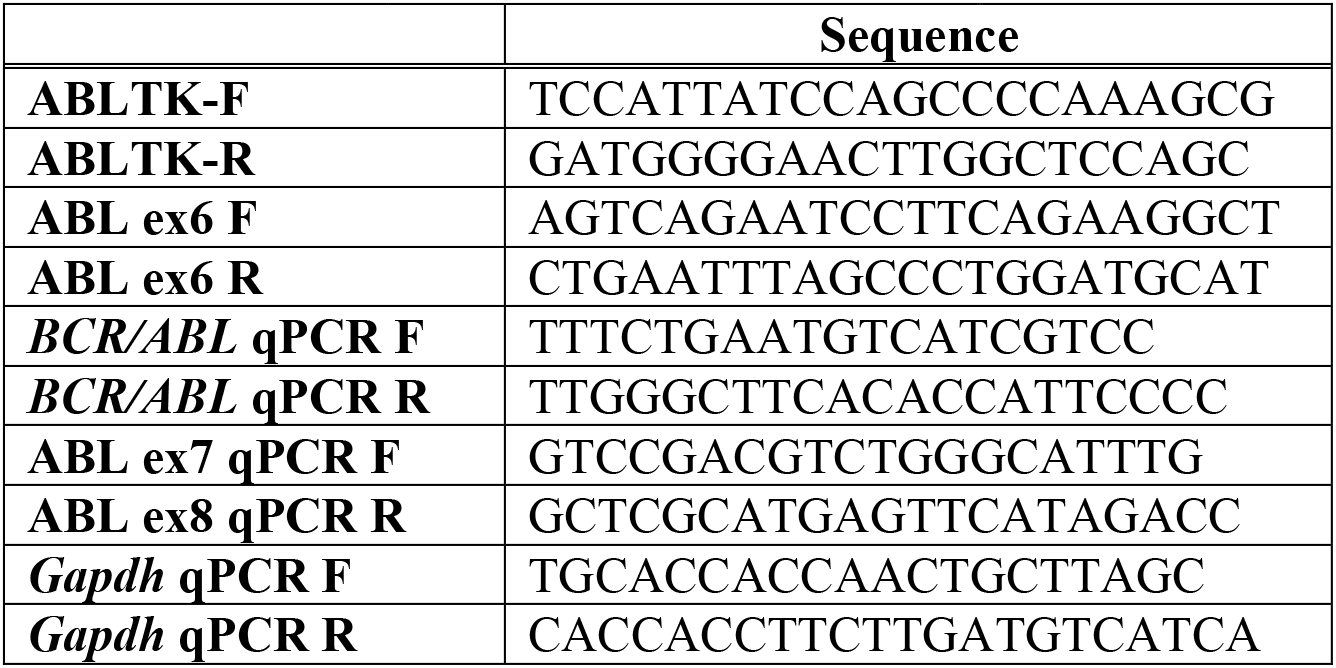
Oligonucleotides

**Supplementary Table 2.**
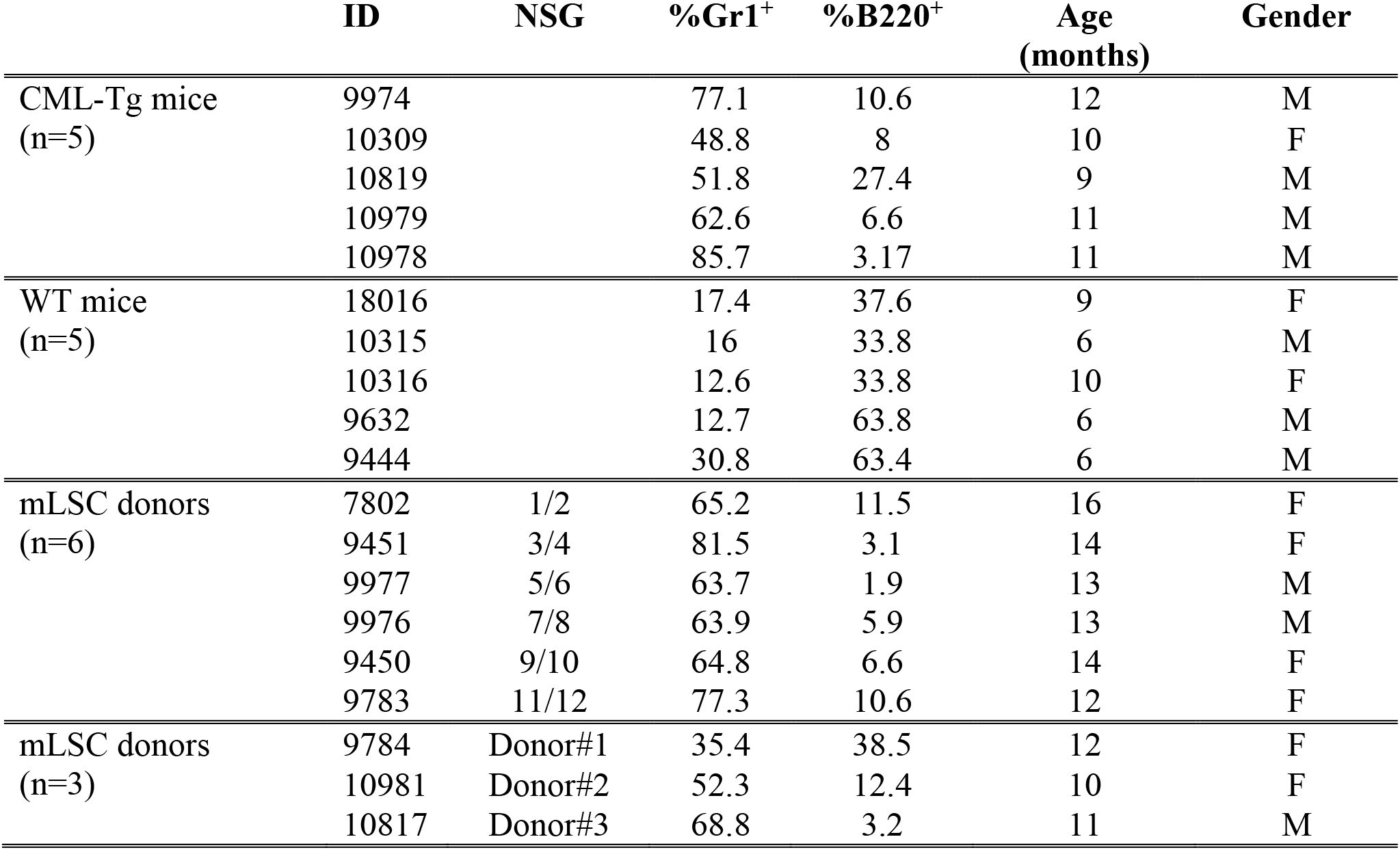
mLSC Donor mice used for mice-mice bone marrow transplantations.

|  | ID | NSG | %Gr1<sup>+</sup> | %B220<sup>+</sup> | Age<br>(months) | Gender |
| --- | --- | --- | --- | --- | --- | --- |
| CML-Tg mice<br>(n=5) | 9974 |  | 77.1 | 10.6 | 12 | M |
|  | 10309 |  | 48.8 | 8 | 10 | F |
|  | 10819 |  | 51.8 | 27.4 | 9 | M |
|  | 10979 |  | 62.6 | 6.6 | 11 | M |
|  | 10978 |  | 85.7 | 3.17 | 11 | M |
| WT mice<br>(n=5) | 18016 |  | 17.4 | 37.6 | 9 | F |
|  | 10315 |  | 16 | 33.8 | 6 | M |
|  | 10316 |  | 12.6 | 33.8 | 10 | F |
|  | 9632 |  | 12.7 | 63.8 | 6 | M |
|  | 9444 |  | 30.8 | 63.4 | 6 | M |
| mLSC donors<br>(n=6) | 7802 | 1/2 | 65.2 | 11.5 | 16 | F |
|  | 9451 | 3/4 | 81.5 | 3.1 | 14 | F |
|  | 9977 | 5/6 | 63.7 | 1.9 | 13 | M |
|  | 9976 | 7/8 | 63.9 | 5.9 | 13 | M |
|  | 9450 | 9/10 | 64.8 | 6.6 | 14 | F |
|  | 9783 | 11/12 | 77.3 | 10.6 | 12 | F |
| mLSC donors<br>(n=3) | 9784 | Donor#1 | 35.4 | 38.5 | 12 | F |
|  | 10981 | Donor#2 | 52.3 | 12.4 | 10 | F |
|  | 10817 | Donor#3 | 68.8 | 3.2 | 11 | M |

**Supplementary Table 3.** FACS Analysis of peripheral blood of NSG mice 120 days after mLSC transplantation.

|  | ID | %Mac1<sup>+</sup> | %Gr1<sup>+</sup> | %Mac1<sup>+</sup>Gr1<sup>+</sup> | %B220<sup>+</sup> | IgM<sup>+</sup> | CD4<sup>+</sup> | CD8<sup>+</sup> |
| --- | --- | --- | --- | --- | --- | --- | --- | --- |
| CML-Tg mice<br>n=5 | 9974 | 51.0 | 77.1 | 61.6 | 10.6 | 5.8 | 2.5 | 2.2 |
|  | 10309 | 65.9 | 48.8 | 42.3 | 8 | 29.7 | 6.3 | 9.5 |
|  | 10819 | 50.7 | 51.8 | 45.8 | 27.4 | 25.1 | 8.9 | 8.4 |
|  | 10979 | 53.0 | 62.6 | 48.2 | 6.6 | 2.9 | 4.9 | 1.8 |
|  | 10978 | 77.0 | 85.7 | 72.8 | 3.17 | 0.7 | 2.8 | 0.9 |
| NSG-Cas9<br>n=6 | 1 | 59.5 | 49.7 | 33.1 | 9.9 | 13.9 | 18.8 | 8.7 |
|  | 3 | 24.8 | 31.6 | 20.1 | 1.0 | 20.0 | 34.5 | 19.4 |
|  | 5 | 64.1 | 73.0 | 44.9 | 2.5 | 1.5 | 0.9 | 0.9 |
|  | 7 | 54.5 | 89.3 | 49.4 | 6.3 | 10.5 | 2.9 | 4.5 |
|  | 9 | 72.9 | 52.3 | 44.0 | 2.0 | 1.5 | 7.9 | 9.1 |
|  | 11 | 63.2 | 67.5 | 40.2 | 4.0 | 1.8 | 2.8 | 1.8 |
| WT mice<br>n=5 | 18016 | 21.8 | 17.4 | 15.9 | 37.6 | 30.7 | 17.8 | 16.3 |
|  | 10315 | 21.6 | 16 | 12.8 | 33.8 | 27.6 | 23.2 | 19.9 |
|  | 10316 | 16.8 | 12.6 | 10.4 | 33.8 | 26.2 | 26.2 | 19.2 |
|  | 9632 | 15.1 | 12.7 | 10.2 | 63.8 | 42.7 | 21.3 | 13.9 |
|  | 9444 | 22.7 | 30.8 | 19.9 | 63.4 | 21.7 | 25.8 | 13.0 |
| NSG-ABL-TK<br>n=6 | 2 | 43.7 | 52.1 | 32.1 | 13.9 | 10.1 | 12.3 | 6.3 |
|  | 4 | 11.9 | 17.9 | 7.2 | 1.0 | 28.8 | 40.9 | 21.2 |
|  | 6 | 38.0 | 35.2 | 22.6 | 27.9 | 13.0 | 11.7 | 5.8 |
|  | 8 | 38.0 | 17.1 | 15.0 | 27.9 | 6.9 | 33.2 | 16.2 |
|  | 10 | 26.9 | 24.0 | 19.0 | 39.5 | 23.6 | 17.9 | 13.9 |
|  | 12 | 29.0 | 26.6 | 14.5 | 33.5 | 12.0 | 19.2 | 15.5 |

**Supplementary Table 4.** FACS Analysis of bone marrow hematological populations in NSG mice 180 days after hLSC transplantation.

|  | **ID** | **%CD34<sup>+</sup>** | **%CD14<sup>+</sup>** | **%CD117<sup>+</sup>** | **%CD19<sup>+</sup>** |
| --- | --- | --- | --- | --- | --- |
| NSG-hLSC CD34<br>n=3 | 13 | 27.0 | 4.6 | 92.7 | 26.5 |
|  | 14 | 12.8 | 3.8 | 62.0 | 22.6 |
|  | 15 | 11.3 | 50.0 | 85.1 | 12.3 |
| NSG- hLSC-Cas9<br>n=7 | 16 | 0.5 | 14.4 | 91.0 | 5.2 |
|  | 17 | 2.9 | 20.6 | 91.9 | 6.6 |
|  | 18 | 23.9 | 34.2 | 85.9 | 20.5 |
|  | 19 | 9.9 | 94.7 | 97.0 | 7.3 |
|  | 20 | 29.6 | 89.5 | 62.0 | 30.4 |
|  | 21 | 18.6 | 64.4 | 90.9 | 12.2 |
|  | 22 | 18.1 | 87.3 | 91.1 | 13.5 |
| NSG-healthy CD34<br>n=3 | 23 | 11.3 | 0.7 | 0.9 | 92.3 |
|  | 24 | 1.8 | 0.9 | 1.6 | 94.9 |
|  | 25 | 3.2 | 1.8 | 2.1 | 91.1 |
| NSG-hLSC-*ABL*-TK<br>n=7 | 26 | 35.6 | 7.8 | 4.3 | 70.5 |
|  | 27 | 1.9 | 16.3 | 9.2 | 84.0 |
|  | 28 | 4.4 | 1.6 | 0.2 | 95.8 |
|  | 29 | 0.5 | 7.2 | 23.7 | 82.4 |
|  | 30 | 0.1 | 0.1 | 0.3 | 95.9 |
|  | 31 | 0.1 | 0 | 55.4 | 80.9 |
|  | 32 | 9.7 | 4.3 | 1.7 | 84.7 |

## Notes

### Competing Interest Statement

The authors have declared no competing interest.

### Summary of Updates

Discussion section has been modified and some bibliographic references has been added.

